# Subtypes of brain change in aging and their associations with cognition and Alzheimer’s disease biomarkers

**DOI:** 10.1101/2024.03.04.583291

**Authors:** Elettra Capogna, Øystein Sørensen, Leiv Otto Watne, James Roe, Marie Strømstad, Ane Victoria Idland, Nathalie Bodd Halaas, Kaj Blennow, Henrik Zetterberg, Kristine Beate Walhovd, the Alzheimer's Disease Neuroimaging Initiative, the Australian Imaging Biomarkers and Lifestyle flagship study of ageing, Anders Martin Fjell, Didac Vidal-Piñeiro

**Affiliations:** Center for Lifespan Changes in Brain and Cognition, Department of Psychology, University of Oslo, 0373 Oslo, Norway; Department of Geriatric Medicine, Akershus University Hospital, Lørenskog, Norway; Institute of Clinical Medicine, Campus Ahus, University of Oslo, Oslo, Norway; Oslo Delirium Research Group, Department of Geriatric Medicine, Oslo University Hospital, Oslo, Norway; Institute of Neuroscience and Physiology, the Sahlgrenska Academy at University of Gothenburg, Mölndal, Sweden; Clinical Neurochemistry Laboratory, Sahlgrenska University Hospital, Mölndal, Sweden; Paris Brain Institute, ICM, Pitié-Salpêtrière Hospital, Sorbonne University, Paris, France; Neurodegenerative Disorder Research Center, Division of Life Sciences and Medicine, and Department of Neurology, Institute on Aging and Brain Disorders, University of Science and Technology of China and First Affiliated Hospital of USTC, Hefei, P.R. China; Department of Neurodegenerative Disease, UCL Institute of Neurology, London, UK; UK Dementia Research Institute at UCL, London, UK; Hong Center for Neurodegenerative Diseases, Hong Kong, China; Wisconsin Alzheimer’s Disease Research Center, University of Wisconsin School of Medicine and Public Health, University of Wisconsin-Madison, Madison, WI, USA; Computational Radiology and Artificial Intelligence, Department of Radiology and Nuclear Medicine, Oslo University Hospital, Oslo, Norway; Institute of Clinical Medicine, Campus Ullevål, University of Oslo, Oslo, Norway

**Keywords:** Memory, Ageotypes, Longitudinal MRI, Cognitively unimpaired older adults, CSF AD biomarkers

## Abstract

Structural brain changes underly cognitive changes in older age and contribute to inter-individual variability in cognition. Here, we assessed how changes in cortical thickness, surface area, and subcortical volume, are related to cognitive change in cognitively unimpaired older adults using structural magnetic resonance imaging (MRI) data-driven clustering. Specifically, we tested (1) which brain structural changes over time predict cognitive change in older age (2) whether these are associated with core cerebrospinal fluid (CSF) Alzheimer’s disease (AD) biomarkers phosphorylated tau (p-tau) and amyloid-β (Aβ42), and (3) the degree of overlap between clusters derived from different structural features. In total 1899 cognitively healthy older adults (50 - 93 years) were followed up to 16 years with neuropsychological and structural MRI assessments, a subsample of which (n = 612) had CSF p-tau and Aβ42 measurements. We applied Monte-Carlo Reference-based Consensus clustering to identify subgroups of older adults based on structural brain change patterns over time. Four clusters for each brain feature were identified, representing the degree of longitudinal brain decline. Each brain feature provided a unique contribution to brain aging as clusters were largely independent across modalities. Cognitive change and baseline cognition were best predicted by cortical area change, whereas higher levels of p-tau and Aβ42 were associated with changes in subcortical volume. These results provide insights into the link between changes in brain morphology and cognition, which may translate to a better understanding of different aging trajectories.

## 1. Introduction

Cognitive changes in healthy aging are partly explained by age-related macrostructural brain changes that may be quantified using repeated structural magnetic resonance imaging (MRI) assessments (Fjell and Walhovd, 2010). However, the extent of age-related changes in brain and cognition differs among older individuals (Lindenberger, 2014), and such differences may be partly underpinned by different patterns of brain aging. The association between brain changes and cognitive change can be assessed by examining different features of morphometric changes, such as cortical thickness, surface area, and subcortical volume. These features have been associated with different aspects of cognition in aging (Nyberg et al., 2023), and with different cerebrospinal fluid (CSF) biomarkers of Alzheimer’s disease (AD) pathology (Fjell et al., 2010; Pettigrew et al., 2016; Wang et al., 2015). MRI data-driven clustering approaches have proven useful for separating subgroups of healthy older participants (“ageotypes”) with different biological, cognitive, and sociodemographic characteristics (Ahadi et al., 2020; Cox et al., 2021). Hence, in the present study, we applied a Monte-Carlo Reference-based consensus clustering algorithm (John et al., 2020) on longitudinal MRI brain features to identify subgroups of cognitively unimpaired older adults based on different structural brain change patterns over time. Moreover, we tested whether the different features of brain change were associated with cognitive changes and with core AD CSF biomarkers the 42 amino acid-long form of amyloid-β (Aβ42) and phosphorylated tau 181 (p-tau) to gain insight into whether brain changes in normal aging can potentially be explained by the presence of AD biomarkers.

In the aging context, longitudinal studies are necessary to capture inter-individual variability in structural brain changes (slope differences), because cross-sectional studies cannot separate aging-specific effects from earlier individual differences (intercept differences) in brain structural measures (Fjell et al., 2014a; Vidal-Piñeiro et al., 2022). This is also supported by a study that found that the underlying factor structure for intercepts versus slopes across brain regions was different, and the correlation patterns between cortical volumetric change were stronger than those observed at baseline in cross-sectional analysis (Cox et al., 2021). Multiple timepoints and long follow-up times are critical to estimate the association between changes in the brain and cognition (Raz and Lindenberger, 2011), and to better understand the neurobiological mechanisms underlying specific cognitive aging processes (Cox et al., 2021; Fjell et al., 2014a).

The use of data-driven clustering, whether based on MRI or cognitive data, is beneficial in assessing the heterogeneity of changes in older participants. This approach was used by Josefsson et al. (2012) who identified ageotypes based on longitudinal trajectories of memory change over 15 years. Participants were divided into maintainers, decliners, and those showing average changes associated with age. Specific environmental and genetic characteristics (such as sex, variance in occupation, education, and physical activity) were related to each group.

Different morphometric features, such as cortical thickness, surface area, and subcortical volume, have been studied to describe inter and intra-individual variation in brain structures. The different features are thought to be largely unrelated to each other (Lemaitre et al., 2012) or even to be negatively associated, as in Storsve and colleagues (2014) where less decrements in cortical area were associated with more cortical thinning. Cortical thickness, cortical area, and subcortical volume decline in aging (Borgeest et al., 2021; Nyberg et al., 2023; Storsve et al., 2014); yet few studies have assessed longitudinal brain changes, taking into account the different brain features (Borgeest et al., 2021; Nyberg et al., 2023; Sele et al., 2021; Storsve et al., 2014). Indeed, cortical area and thickness seem to reflect distinct underlying neurobiological mechanisms that are differently affected in aging (Storsve et al., 2014), show specific regional changes, and have a negative genetic correlation (Grasby et al., 2020). Assessing change in the different features of brain aging independently and considering to which degree they complement each other may translate to a better understanding of aging brain heterogeneity.

So far, there is inconclusive evidence regarding the association between structural brain changes and cognitive changes in aging, and most of the evidence is based on cross-sectional studies, with few exeptions. Thickness, but not area, changes were related to fluid intelligence changes (Sele et al., 2021) and memory changes, especially in the medial temporal lobe, as described in one study (Fjell et al., 2014b). Other studies found that surface area changes were associated with changes in proxy measures of fluid intelligence (Borgeest et al., 2021). Nyberg et al. (2023) described a significant association of surface area changes with a speed of processing test. Another study found positive associations between a general cognitive ability (GCA) factor and brain features, but with different results: indeed, higher baseline GCA was associated with greater cortical area at baseline and less cortical thinning over time (Walhovd et al., 2022). Finally, there is a general agreement in the literature regarding the positive association between hippocampal volume loss and episodic memory decline (Capogna et al., 2023a; Gorbach et al., 2020, 2017; Persson et al., 2012).

Decreased CSF Aβ42 (reflecting amyloid accumulation in the brain tissue) and increased p-tau (reflecting a neuronal response to Aβ pathology) concentrations are considered two of the key biomarker hallmarks of AD pathology (Jack et al., 2018), and their changes are identified in the early stage of the AD continuum, without the presence of any cognitive symptoms. Hence, it is relevant to understand the relationship between these biomarkers and the different structural features of brain change in cognitively unimpaired older adults, to better identify how these brain changes may be explained by the presence of AD biomarkers along an aging-disease continuum. CSF Aβ42 and p-tau biomarker changes have been associated with lower cortical thickness and subcortical volume atrophy in AD-vulnerable regions in cognitively unimpaired older adults (Arenaza-Urquijo et al., 2013; Pettigrew et al., 2016; Wang et al., 2015). However, the relationship between CSF Aβ42 and brain atrophy is inconclusive (Fjell et al., 2014a), whereas p-tau shows a stronger association with medial temporal lobe (MTL) atrophy, following the time course of cognitive decline (Pettigrew et al., 2017; Vidal-Piñeiro et al., 2022; Wisse et al., 2022). To our knowledge, no study has explored the relationship between CSF core AD biomarkers and surface area changes. Moreover, early CSF Aβ42 and p-tau biomarker changes (notably for p-tau, unclear for Aβ42 (Parent et al., 2023)) have been described as predictive of future cognitive decline, especially in episodic memory, although the overall effects were small (Clark et al., 2018; Hedden et al., 2013; Stomrud et al., 2007).

In the present study, we investigated the inter-individual patterns of structural brain changes in normal aging using a consensus clustering algorithm. This approach allowed us to identify subgroups, i.e. clusters, among older participants. We used different indices of brain morphology, namely cortical thickness, cortical area, and subcortical volume to better understand the degree to which each modality contributes independently to brain longitudinal decline, and the degree of overlap across them. Moreover, we assessed whether the clusters were related to different trajectories of episodic memory function and global cognition, as measured by dementia screening tools. Finally, we tested the associations between brain changes and baseline concentrations of core AD CSF biomarkers (Aβ42 and p-tau, as well as the p-tau/Aβ42 ratio).

## 2 Material and methods

### 2.1 Participants

The total sample included 1899 cognitively healthy older participants (1080 females, mean age = 69.88 years, standard deviation [SD] = 7.90, age range = 50.11 – 93.01 years) from 7 cohorts: COGNORM (Idland et al., 2017), the Alzheimer’s Disease Neuroimaging Initiative (ADNI) (Mueller et al., 2005), the Open Access Series of Imaging Studies (OASIS3) (LaMontagne et al., 2019), the Australian Imaging, Biomarker & Lifestyle Flagship Study of Ageing (AIBL) (Ellis et al., 2009), the Harvard Aging Brain Study (HABS) (Dagley et al., 2017), the Pre-symptomatic Evaluation of Novel or Experimental Treatments for AD (PREVENT-AD) program (Breitner et al., 2016; Tremblay-Mercier et al., 2021), and the Center for Lifespan Changes in Brain and Cognition (LCBC) dataset (Fjell et al., 2023). Data were collected by previously cited groups. See **Table 1** for more details on each dataset. The common inclusion criteria were as follows: minimum age of 50 years, total follow-up time of at least 1 year, and inclusion of scanners with 15 or more measurements to reduce noise and bias in the analysis. Moreover, the participants were required to be cognitively unimpaired at baseline according to a battery of neuropsychological tests. See specific inclusion criteria for each cohort in Supplementary Information (**SI**). Longitudinal structural MRI scans were available for up to 15.84 years (mean = 4.81 [2.81] years). At baseline, participants showing concurrent mild cognitive impairment, AD, or other severe neurological disorders were excluded from the analysis. All participants provided written informed consent, and the studies were approved by the relevant ethical committees and conducted in accordance with the Declaration of Helsinki.

**Table 1.** Cohort characteristics.

|  | **COGNORM** | **ADNI** | **OASIS 3** | **AIBL** | **HABS** | **PREVENT-**<br>**AD** | **LCBC** | **Total** |
| --- | --- | --- | --- | --- | --- | --- | --- | --- |
| **N (F:M)** | 95 (52:43) | 544 (293:251) | 518 (292:226) | 149 (75:74) | 166 (99:67) | 229 (161:68) | 198 (108:90) | 1899<br>(1080:819) |
| **Mean Age** | 73.40 (6.21) | 73.22 (5.98) | 69.00 (8.41) | 71.09 (6.34) | 72.78 (6.04) | 63.56 (5.03) | 65.23 (9.64) | 69.88 (7.90) |
| **Age range** | 64.74 – 89.79 | 55.80 – 89.90 | 50.11 – 93.01 | 60.00 – 87.00 | 62.50 – 87.75 | 55.13 – 84.22 | 50.39 – 84.47 | 50.11 – 93.01 |
| **Time MRI**<br>**follow-up** | 5.94 (2.61) | 4.12 (2.68) | 5.13 (3.35) | 4.04 (1.62) | 4.88 (0.97) | 2.88 (1.13) | 6.30 (2.80) | 4.81 (2.81) |
| **MRI obs (n)** | 370 (3.90<br>[1.36]) | 2492 (4.58<br>[2.42]) | 1652 (3.19<br>[1.49]) | 498 (3.34<br>[1.10]) | 504 (3.04<br>[0.53]) | 1203 (5.23<br>[1.39]) | 574 (2.90<br>[0.83]) | 7293 (3.84<br>[1.86]) |
| **Global cognition**<br>**obs (n)** | 609 (6.41<br>[0.99]) | 2614 (4.80<br>[2.52]) | 3584 (6.96<br>[3.98]) | 598 (4.01<br>[1.17]) | 988 (5.95<br>[0.21]) | 229 (1 [0]) | 525 (2.65<br>[0.86]) | 9147 (4.54<br>[1.39]) |
| **Memory obs (n)** | 609 (6.41<br>[0.99]) | 2836 (5.21<br>[2.67]) | 493 (4.48<br>[2.68]) | 507 (3.40<br>[1.13]) | 993 (5.98<br>[0.80]) | 969 (4.25<br>[1.39]) | 420 (2.80<br>[0.79]) | 6827 (4.64<br>[1.49]) |
| **Education (years)** | 14.94 (3.97) | 16.50 (2.56) | 15.89 (2.63) | - | 16.14 (3.01) | 15.25 (3.27) | 16.04 (2.77) | 15.97 (2.88) |
| *APOE* ε4 (-/+) | 48:37 | 368:170 | 332:186 | 99:50 | 118:46 | 146:83 | 50:21 | 1161:593 |

Descriptive statistics represent mean (SD). N = number of subjects. Obs = total number of observations. *The MRI obs* row shows the mean ([SD])
of the number of observations per participant and the total number. The same applies to global cognition and memory. Note that the term "global
cognition" specifically pertains to the screening tests outlined below. *APOE* ε4 = non-carriers:carriers. Participants with heterozygous or
homozygous ε4 alleles were regarded as carriers.

### 2.2 MRI acquisition and preprocessing

Structural T1-weighted (T1w) MPRAGE scans were collected using 1.5 and 3 T scanners. See information on scanner parameters and scanners per dataset in the SI. Images were transformed into the Brain Imaging Data Structure (BIDS) format (Gorgolewski et al., 2016). Clinica software was used for the ADNI, AIBL, and HABS BIDS transformations (Routier et al., 2021; Samper-González et al., 2018). We used the longitudinal FreeSurfer v.7.1.0 stream (Reuter et al., 2012) for cortical reconstruction of the structural T1w scans (Dale et al., 1999; Fischl et al., 1999). Briefly, the images were processed using the cross-sectional stream, which includes the removal of nonbrain tissues, Talairach transformation, intensity correction, tissue and volumetric segmentation, cortical surface reconstruction, and cortical parcellation. Next, an unbiased within-subject template space based on all cross-sectional images was created for each participant, using robust, inverse-consistent registration (Reuter et al., 2010). The processing of each time point was then reinitialized with common information from the within-subject template, to increase reliability and statistical power. Data were summarized based on the Destrieux atlas (Destrieux et al., 2010) for cortical thickness and cortical area measures (74 features) and the *aseg* atlas for subcortical volumetric data (17 features) (Fischl et al., 2002).

### 2.3 Computation of intercept and slope measures for ROIs per participant

We focused on three indices of cerebral morphology: cortical thickness, surface area, and subcortical volume. For each region of interest (ROI), we regressed out the effects of mean age across timepoints for each participant using generalized additive mixed models (GAMM) (Wood, 2017), as implemented in the *gamm4* package. Age was introduced as a smooth term, while random intercepts were included for each dataset, scanner, and participant. To compute the slope of change for each participant, we fit a linear regression model for each participant and ROI with the GAMM model residuals (as dependent variable) and time equal to the difference between age at a given observation and the individual’s mean age. Participants without longitudinal MRI data and those with follow-up intervals of < 1 year were excluded from further analysis and were not included in the final sample. Next, we replaced outlier values (> ±5 SD from the mean) using the *mice* package (Buuren and Groothuis-Oudshoorn, 2011) (0.003% of observation values were replaced). The final output yielded a total of 330 structural MRI features (148 cortical thickness, 148 surface area, and 34 subcortical volumetric bilateral ROIs that contained slope data). Finally, the values were scaled for each feature.

### 2.4 Consensus clustering of brain data

Slope data were clustered based on the M3C clustering algorithm – a Monte-Carlo Reference-based Consensus algorithm - as implemented in the *M3C* package (John et al., 2020). Consensus algorithms (Monti et al., 2003) are based on the idea that the ideal cluster should be stable despite resampling, that is, that individuals should always or never be clustered together in the face of iterative resampling. Such methods have gained popularity as they produce more robust results, reduce bias, and provide estimates of the error.

An important challenge in consensus clustering is selecting the number of clusters (K). The most popular criteria either require subjective decisions or show biases towards small or high K-solutions. Furthermore, most approaches cannot test whether the desired solution is better than K = 1 (i.e., that the data comes from a single distribution). M3C solves both problems by generating Monte Carlo simulations that preserve the covariance structure. M3C provides null stability scores for a range of K values, which are then compared with real-data solutions. Here, we used the Relative Cluster Stability Index (RCSI) metric to select K based on the proportion of ambiguous clusters (PAC) scores. RCSI p-values were further derived to test the null hypothesis of K = 1 at each value of K. We used a spectral clustering algorithm (Ng et al., 2001) as it is capable of coping with complex data structures. The remaining parameters were set to the default values of *M3C* version 1.24.0. The algorithm was separately applied to each of the three morphometric brain measures. To explore the data structure, we projected the cluster outcome (i.e., individual assignments) onto main components of brain change (i.e., components capturing the main axis of variability of brain data) (n = 4) and carried out an ANOVA using cluster assignment as the factor of interest. When significant (Bonferroni-corrected), post-hoc pairwise comparisons were performed. Finally, for each cluster and feature, we estimated the mean values.

### 2.5 Degree of overlap between cluster solutions

We carried out an analysis to establish whether the different structural modalities were statistically related to each other, i.e., whether participants belonged to different clusters or the same cluster across the various morphometric brain features. The explorative analysis indicated that clustering was based on one principal component from a Principal Component Analysis (PCA), and all the clustering solutions resulted in four groups. See **Supplementary Figure 1**. Thus, for clarity, we renamed the clusters based on their mean ROIs change values as follows: Decline, Mild Decline, Mild Maintenance, Maintenance. Cohen’s kappa was used to assess the agreement among clusters, as implemented in the *psych* and *irr* R-packages. A weighted kappa coefficient was applied because of the ordinal characteristics of the clusters and to stress the large discrepancies in ratings more than the small ones (Sim and Wright, 2005).

### 2.6 Cognitive functions over time

We focused on memory and global cognitive impairment because of their relevance in aging. For the global cognitive impairment factor, we used the longitudinal scores from the Mini-Mental State Examination (MMSE) (Folstein et al., 1975) for all samples except PREVENT-AD, for which the Montreal Cognitive Assessment (MOCA) was used (Nasreddine et al., 2005). We used these screening tests as a global measure of cognitive function (Garcia-Diaz et al., 2014; Matsushima et al., 2015), acknowledging their sensitivity to dementia and cognitive decline in the aging-disease continuum. The number of participants included was n = 1896. Within cohorts, we scaled the longitudinal scores based on the mean and SD at the first time point (same procedure applied below for the memory scores). Note that PREVENT-AD did not include longitudinal MOCA scores. Moving forward, we will collectively refer to the output of these screening tests as ‘global cognition’, as they are both sensitive to premorbid global cognitive decline. For the memory factor, we selected the precomputed ADNI-MEM (Crane et al., 2012) for the ADNI dataset. For the other datasets, we used the Immediate and Delay scores in the Word List Memory Task (CERAD) (Morris et al., 1989) for COGNORM, and the short delay and delayed score of the Logical Memory Test (Wechsler, 1987) for AIBL, HABS, and OASIS3. For PREVENT-AD, we used the memory index score (Immediate and Delayed) obtained from the Repeatable Battery for Assessment of Neuropsychological Status (RBANS) (Randolph et al., 1998; Tremblay-Mercier et al., 2021) and the short delay, delayed, and total learning from the California Verbal Learning Test (Delis et al., 2000) for LCBC. Then we performed separate PCA on the first timepoint in each dataset with multiple memory variables. The loadings for the first component were used to calculate scores for the first principal component across all timepoints (Capogna et al., 2023b). The *prcomp* function was used for the PCA. Furthermore, for both memory and global cognition factors, we regressed the effects of age using GAMMs (Wood, 2017). Age was introduced as a smooth term and a test-retest variable as a dichotomic covariate to account for training effects (Capogna et al., 2023b), and random intercepts were included for each participant in the model. To compute the slope, we first extracted the residuals from the GAMMs, and then we ran a linear regression model for each participant with age as the predictor and residuals as outcome. For the global cognition factor, longitudinal results were available for 1649 participants, and the memory change factor for 1442 participants.

### 2.7 CSF collection, analysis and computation of intercept and slope

CSF data were available for three cohorts: ADNI, COGNORM, and PREVENT AD (total number of participants available = 612). For ADNI, CSF Aβ42 and p-tau concentrations were measured using Elecsys phosphorylated-tau 181 (p-tau) and β-amyloid (Aβ42) CSF immunoassays (*UPENNBIOMK9.csv* ADNI file). CSF collection for COGNORM has been thoroughly described previously (Idland et al., 2017). Briefly, CSF samples were analyzed at the Clinical Neurochemistry Laboratory of Sahlgrenska University Hospital (Mölndal, Sweden). CSF concentrations of Aβ42 and p-tau were measured using the INNOTEST enzyme-linked immunosorbent assay (ELISA; Fujirebio, Ghent, Belgium). CSF collection for PREVENT-AD has been described previously (Tremblay-Mercier et al., 2021). CSF samples for Aβ42 and p-tau 181 were measured using an INNOTEST enzyme-linked immunosorbent assay. We had 608 and 611 cross-sectional values for p-tau and Aβ42 respectively, and longitudinal values available for 327 and 328 participants for p-tau and Aβ42, respectively. Within each cohort, we first scaled each CSF value based on the mean and SD at the first timepoint. To compute the intercept and slope measure for CSF biomarkers, we fitted a linear regression model for each participant with the CSF scaled value as the dependent variable and time equal to the difference between age at a given observation and age at baseline. Due to the relatively small number of participants with longitudinal CSF data, the longitudinal biomarkers analyses are deemed exploratory (see **Supplementary Table 3**). See **Table 2** for descriptive CSF data.

**Table 2.** Cross-sectional and longitudinal info CSF AD biomarkers.

|  | ADNI | PREVENT-AD | COGNORM |
| --- | --- | --- | --- |
| **N** | 412 | 106 | 94 |
| **CSF p-tau bsl** (pg/mL) | 21.51 (8.74) | 48.16 (17.67) | 61.40 (18.96) |
| **CSF Aβ42 bsl** (pg/mL) | 1359.10 (649.67) | 1152.40 (270.77) | 729.48 (205.77) |
| **Time from first MRI** (years) | 1.47 (2.24) | 1.43 (1.33) | 0 (0) |
| **Interval follow- up** (range in years) | -3.74– 10.28 | 0.22 – 4.58 | 2.88 – 5.69 |
| **N follow-up** | 221 | 76 | 34 |
| **CSF p-tau total obs** | 802 | 351 | 66 |
| **CSF Aβ42 total obs** | 809 | 350 | 66 |

N = number of participants (with MRI available for clustering) with AD biomarkers available.
Bsl = baseline value. The Time variable represents the mean (SD) of years between CSF collections and baseline first MRI measurement. The interval follow-up refers to the range of Time (see above) in years of CSF longitudinal collections, excluding the first CSF assessment.
N follow-up represents the number of participants with at least 2 CSF measurements over time.
Obs = number of total observations for each CSF biomarker of interest.

### 2.8 Statistical Analysis

All analyses were performed in the R environment (R Core Team, 2022). A chi-square test was used to assess whether the clusters were associated with specific socio-demographic variables such as Sex, *APOE* ε4, and Cohort variables. We used the *chisq.posthoc.test* package (Beasley and Schumacker, 1995) to assess the cluster driving the significant associations. Linear mixed-effects models (LME), as implemented in the *lme4* R-package (Bates et al., 2015), were used to assess whether the cluster assignments differed in education and mean age levels. Moreover, we used LME to compute the effect of cluster assignment on memory and global cognition intercept and change. Sex and mean age were introduced as covariates of no interest. Random intercepts per cohort were also included. In addition, 4-group ANOVA models were run on the outputs of the LME models. The models were corrected for multiple comparisons using the false discovery rate and Benjamini-Hochberg correction (pFDR) (Benjamini and Hochberg, 1995). Specifically, we corrected the p-values from all the models separately for each dependent domain (memory, global cognition, and CSF AD biomarkers). If the output was significant, we applied multiple comparisons of means, as implemented in the *multcomp* R-package (Westfall, 2010), that displayed the adjusted p-values, using a single-step method. The same procedure described above was run in a subsample (n = 612) to assess the relationship of brain change clusters with the CSF AD biomarkers. In the p-tau model, we also included baseline CSF Aβ42 as a covariate. We also tested the association with the p-tau/Aβ42 ratio.

### 2.9 Automated model selection

We tested the combined effects of cluster assignments (for changes in thickness, area, and subcortical volume) and their interactions on explaining memory, global cognition, CSF AD biomarkers, intercept, and slope (except for core CSF AD biomarkers). We used a LASSO algorithm (Tibshirani, 1996), that performs a variable selection to maximize the prediction accuracy, as implemented in the *gglasso* package (Yang and Zou, 2015) to automatically identify the best-performing model to explain the cognitive and biomarker changes. We used grouped LASSO to model the categorical properties of clusters, that is, both the different regressors for the main effects of clusters and their interactions were grouped, so the outcome either provided coefficients for all the conditions or none. First, we created a matrix of predictors (X) containing the main effects and all the interactions among clusters, while we set Sex and mean Age as fixed variables. We defined the response (y) as the cognitive or AD biomarkers of interest in prediction. We applied the function *cv.gglasso*, employing 10-folds cross-validation to determine the optimal smoothing λ parameter. We report the results at two different λ: λ at minimum RMSE, and at the largest value of λ within 1 standard error of λ minimum which leads to more conservative results.

## 3. Results

### 3.1 Clustering solutions for brain features and mean values of each cluster

We identified 4 clusters for each brain feature of interest. See **Figure 1** for a visual representation of the results and SI in [Zenodo] at https://doi.org/10.5281/zenodo.10365469 for the stats of the three features. The PCA and the visual exploration of the results suggested that clusters were defined based on a main axis (component) of decline. See **Supplementary Figure 1**. We reordered the clusters from those showing a steeper overall decline to those that displayed – comparatively – less decline. For cortical thickness change, we found a high effect of bilateral temporal and inferior parietal regions on cluster assignment. To some degree, we observed a similar pattern for surface area changes, although weaker and more prominent in the left superior frontal and temporal regions. Subcortical volume cluster assignment was especially influenced by hippocampus decline and ventricular expansion. Henceforth, the clusters are renamed as decline, mild decline, mild maintenance, and maintenance. See **Figures 2, 3**, and **4** for a visual representation of the differences between the different clusters in each analysis.

**Figure 1.**
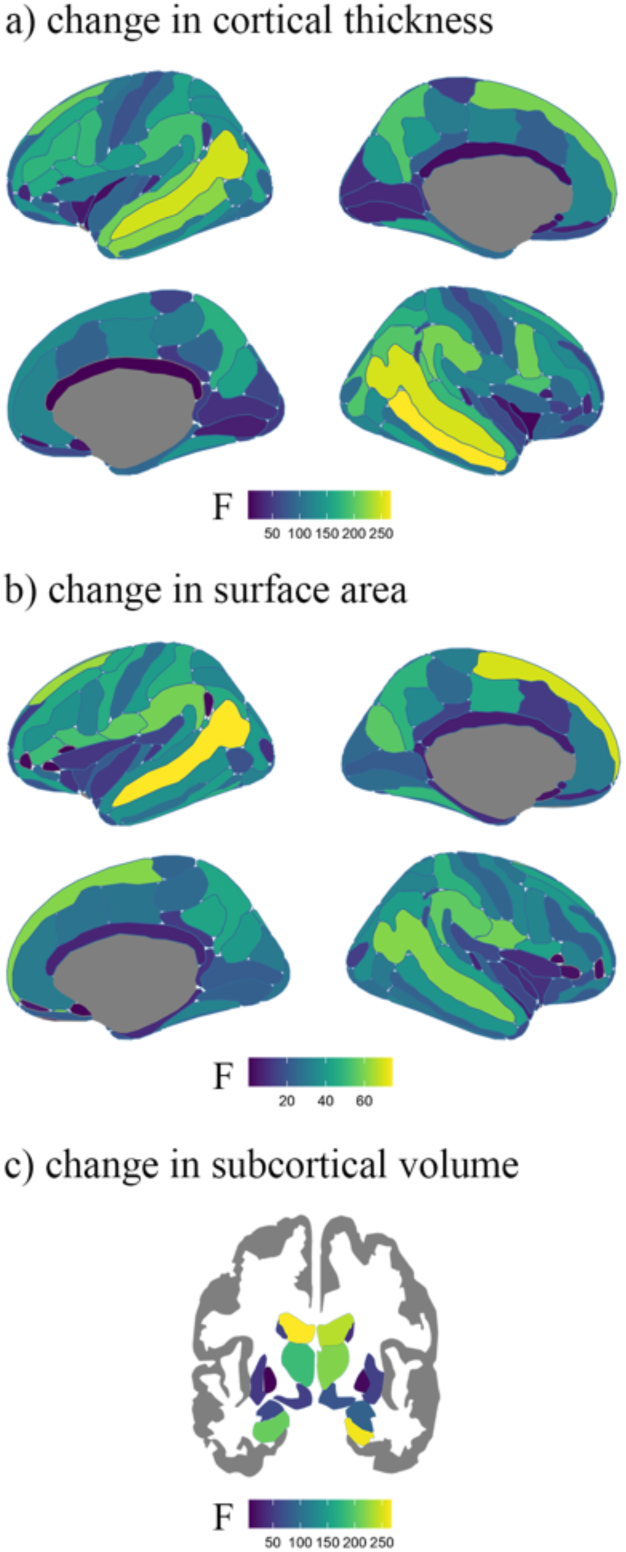
ANOVA output of cluster assignment. ANOVA output of brain change respectively for cluster assignments of cortical thickness, surface area, and subcortical volume. The F-values represent the influence of each region in the cluster assignment. Yellow regions represent more importance and blue regions represent less importance. ROIs were based on the Destrieux atlas (Destrieux et al., 2010) for cortical thickness and cortical area, and the *aseg* atlas (Fischl et al., 2002) for subcortical volumetric data.

**Figure 2.**
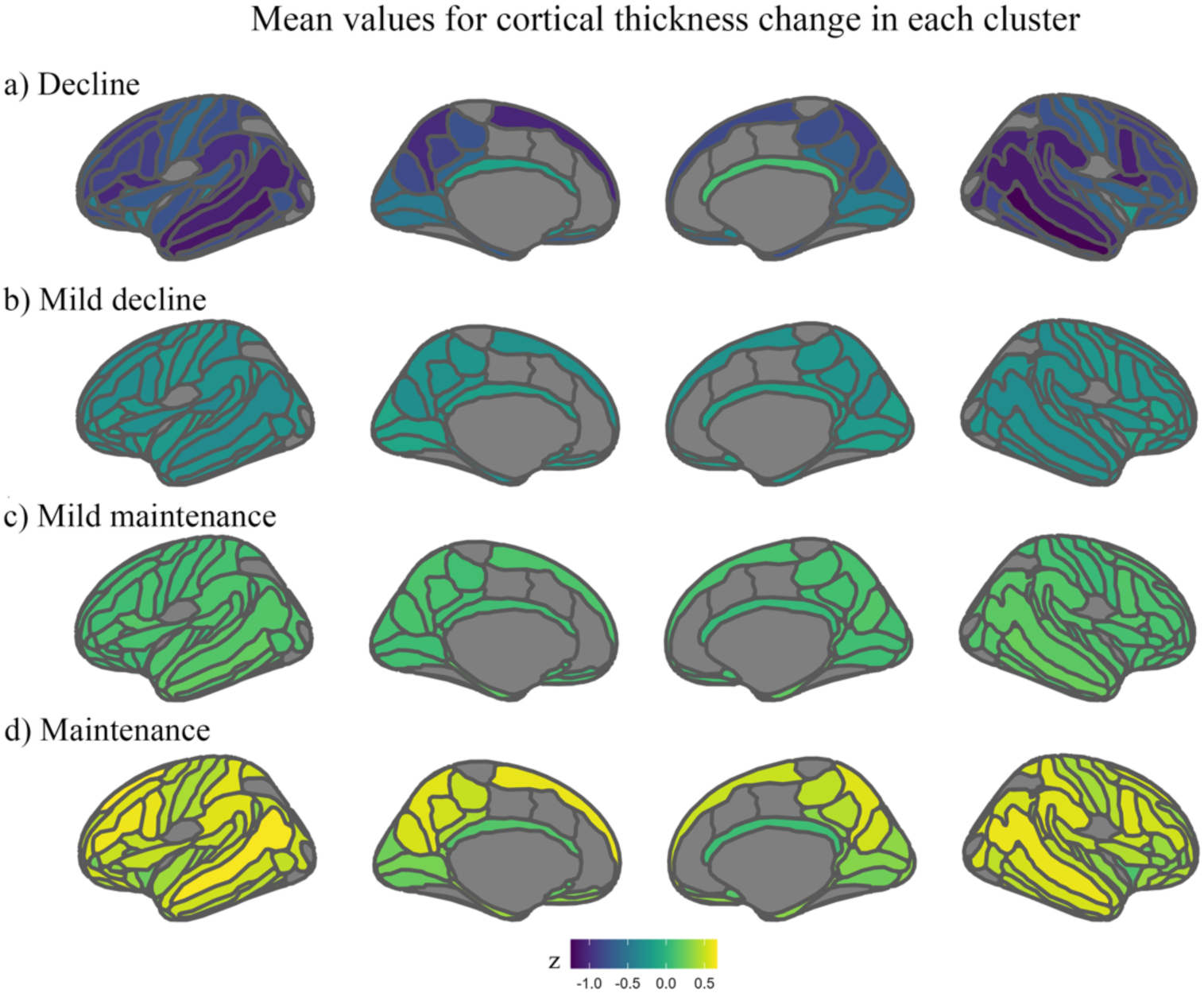
Mean values for cortical thickness change in each cluster. Mean thickness change (z scores) values for each of the four clusters. Yellow represents more positive values and less change in thickness over time, while indigo represents more negative values and more thinning over time. A) decline cluster; b) mild decline cluster; c) mild maintenance cluster; d) maintenance cluster. ROIs were based on the Destrieux atlas (Destrieux et al., 2010) for cortical thickness. ROIs without an overlay are not significant (pFDR < 0.05).

**Figure 3.**
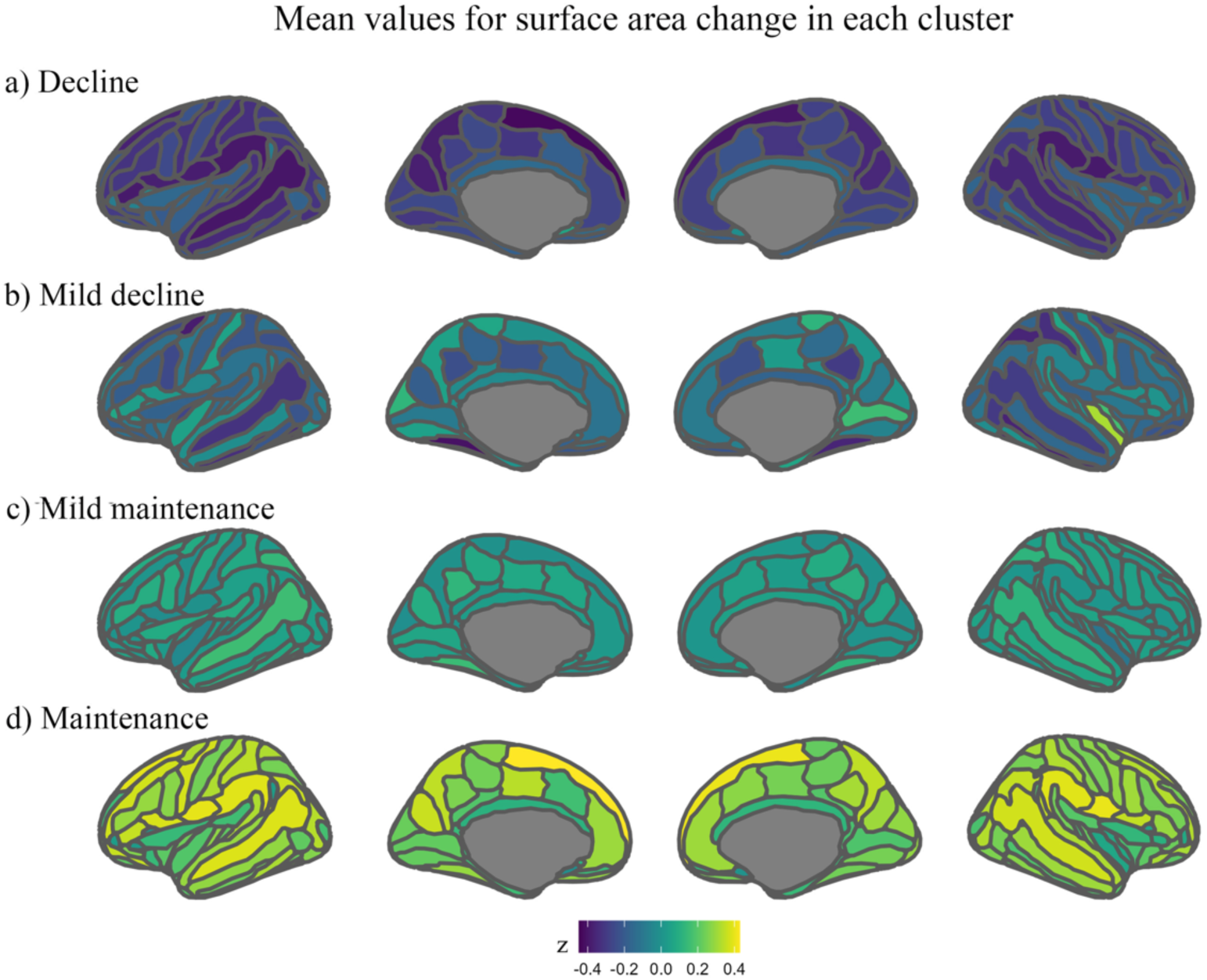
Mean values for surface area change in each cluster. Mean area change (z scores) values for each of the four clusters. Yellow represents more positive values and less change in area over time, while indigo represents more negative values and more change in area over time. A) decline cluster; b) mild decline cluster; c) mild maintenance cluster; d) maintenance cluster. ROIs were based on the Destrieux atlas (Destrieux et al., 2010) for surface area. ROIs without an overlay are not significant (pFDR < 0.05).

**Figure 4.**
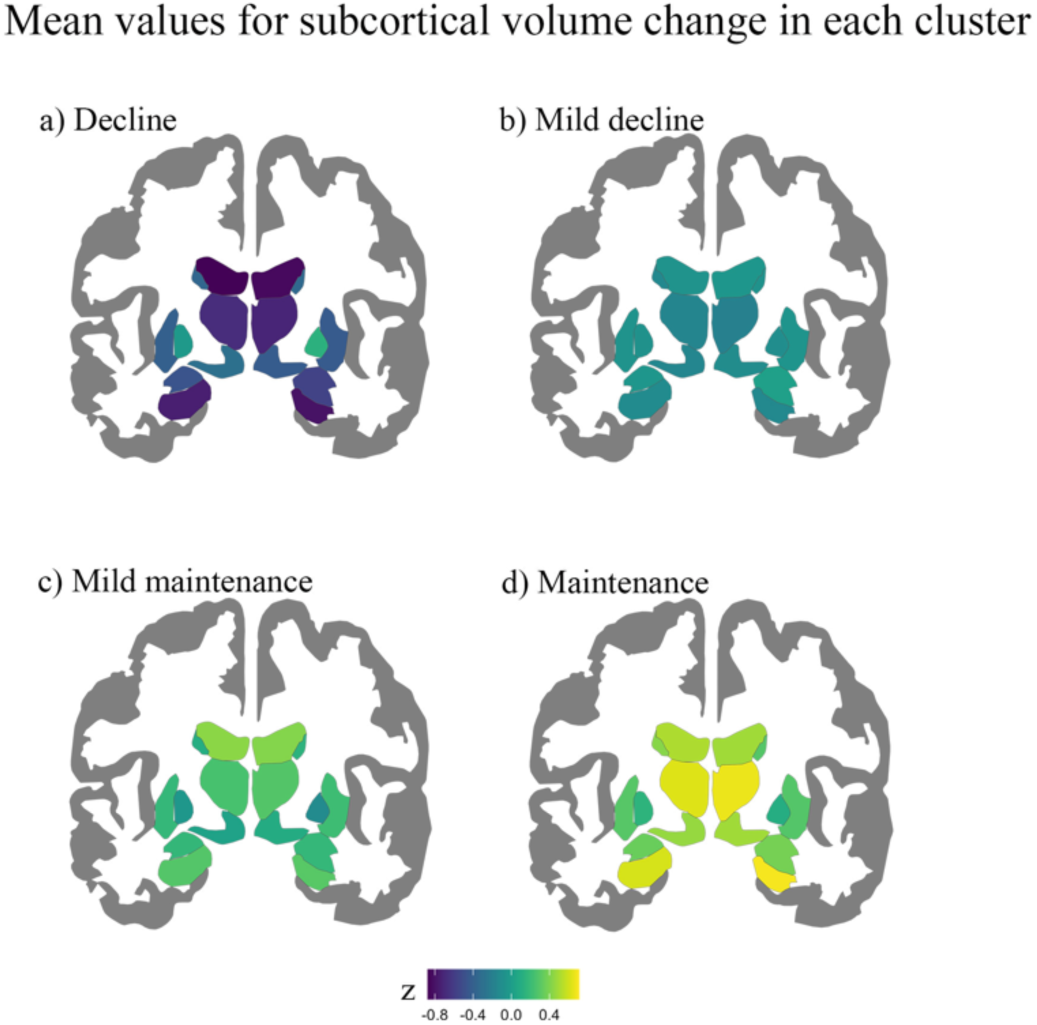
Mean values for subcortical volume change in each cluster. Mean subcortical change (z scores) values for each of the four clusters. Yellow represents more positive values and less change in subcortical volume, while indigo represents more negative values and more subcortical volume decline. A) decline cluster; b) mild decline cluster; c) mild maintenance cluster; d) maintenance cluster. ROIs were based on the *aseg* atlas (Fischl et al., 2002). ROIs without an overlay are not significant (pFDR < 0.05).

### 3.2 Degree of overlap between brain features

We next tested whether participants belonged to different clusters or the same cluster across the various morphometric brain measures. The cluster assignment for each brain feature is summarized in **Figure 5**. The weighted Cohen’s kappa coefficient for correspondence in cluster assignment for thickness and area is κ = 0.08 (p < 0.001), which means that the agreement between clustering of different modalities was slight (as per Landis and Koch, 1977). This suggests, as also previously reported, that thickness and surface area are two largely unrelated and independent morphometric characteristics of aging (Storsve et al., 2014). The agreement between participants being classified on the same clusters for subcortical volume and thickness is weighted κ = 0.29 (p < 001), often interpreted as “fair” (Landis and Koch, 1977), whereas the weighted Cohen’s kappa coefficient for subcortical volume and area is κ = 0.19 (p < 0.001).

**Figure 5.**
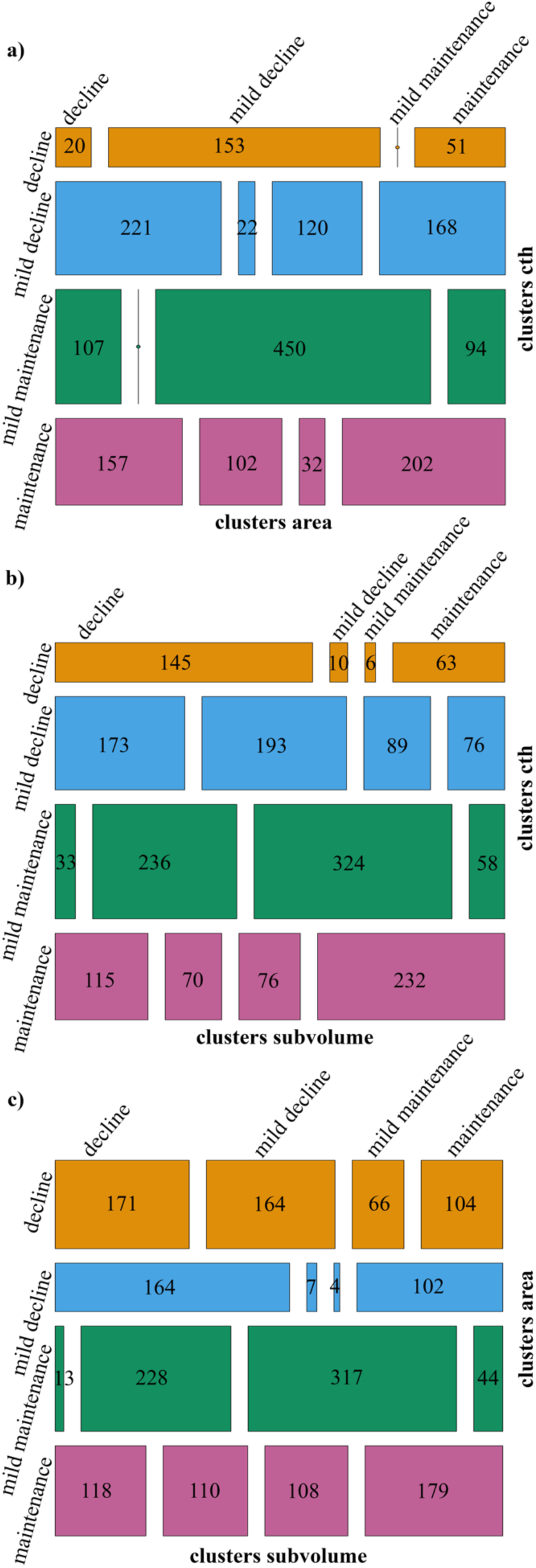
Contingency tables for brain cluster assignment. Mosaic plots reflecting the output of the contingency tables (*vcd R* package). The dimensions of each box are proportional to the number of participants grouped into the clusters based on the different structural modality. The kappa value is calculated based on the discrepancy between the diagonal boxes and those that are not located on the diagonal (agreement vs. disagreement). Cells without a number mean 0 participants belonged to the two different clusters. A) Table for clusters based on thickness change versus clusters based on area change; b) table for clusters based on thickness change versus clusters based on subcortical volume change; c) table for clusters based on area change versus clusters based on subcortical volume change.

### 3.3 Associations between brain cluster assignment and genetic-environmental variables

We then assessed whether the cluster assignment (for brain feature) differed for sex, age, education, *APOE* ε4 status, and cohort. No associations were found with education level. Changes in cortical thickness (cluster assignment) were associated with age (F = 10.97, df1 = 1888.8, df2 = 3, pFDR < 0.001), and *APOE* ε4 status (χ^2^ = 11.86, df residual = 1751, pFDR = 0.01). Changes in cortical surface clusters were related to age (F = 32.56, df1 = 1889, df2 = 3, pFDR < 0.001), sex (χ^2^ = 41.44, df residual = 1896, pFDR < 0.001), and *APOE* ε4 status (χ^2^ = 15.34, df residual = 1751, pFDR < 0.01). Changes in subcortical volume clusters were also related to age (F = 19.98, df1 = 1889.5, df2 = 3, pFDR < 0.001), sex (χ^2^ = 52.15, df residuals = 1896, pFDR < 0.001), and *APOE* ε4 status (χ^2^ = 19.53, df residual = 1751, pFDR < 0.001). Overall, the ANOVA results were in the expected direction, with clusters showing relative brain maintenance having lower age, lower representation of *APOE* ε4 carriers, and less males, whereas clusters showing more brain decline had higher age and a higher representation of *APOE* ε4 carriers and males. See **Supplementary Table 1** for the direction of the significant post-hoc associations between cluster assignment and these genetic and environmental variables.

### 3.4 Associations between brain cluster assignment and cognitive functions

We then assessed the relationship between cluster assignment, intercept and change in memory and global cognition, using LME and 4-group ANOVA models. The results are presented in **Table 3** (including the post-hoc multiple comparisons), for a visual representation see **Figure 6**. We found significant associations between global cognition intercept and changes in cortical area (F = 15.69, df1 = 1889, df2 = 3, pFDR < 0.001), thickness (F = 16.21, df1 = 1889, df2 = 3, pFDR < 0.001), and subcortical volume (F = 15.88, df1 = 1889, df2 = 3, pFDR < 0.001). The ANOVA results were in the expected direction, with clusters showing relative brain maintenance displaying higher cognition, and those showing more brain decline exhibiting lower cognition. See the post-hoc comparisons across groups, that is, which specific clusters had significantly different values in **Table 3** and **Supplementary Table 2**. Global cognition changes were associated with surface area changes (cluster assignment) over time (F = 4.16, df1 = 1605.9, df2 = 3, pFDR = 0.009, post-hoc: lower cognition for mild decline cluster). No significant relationship between cluster assignment and memory change and intercept survived correction for multiple comparisons; all the results were above pFDR > 0.05.

**Figure 6.**
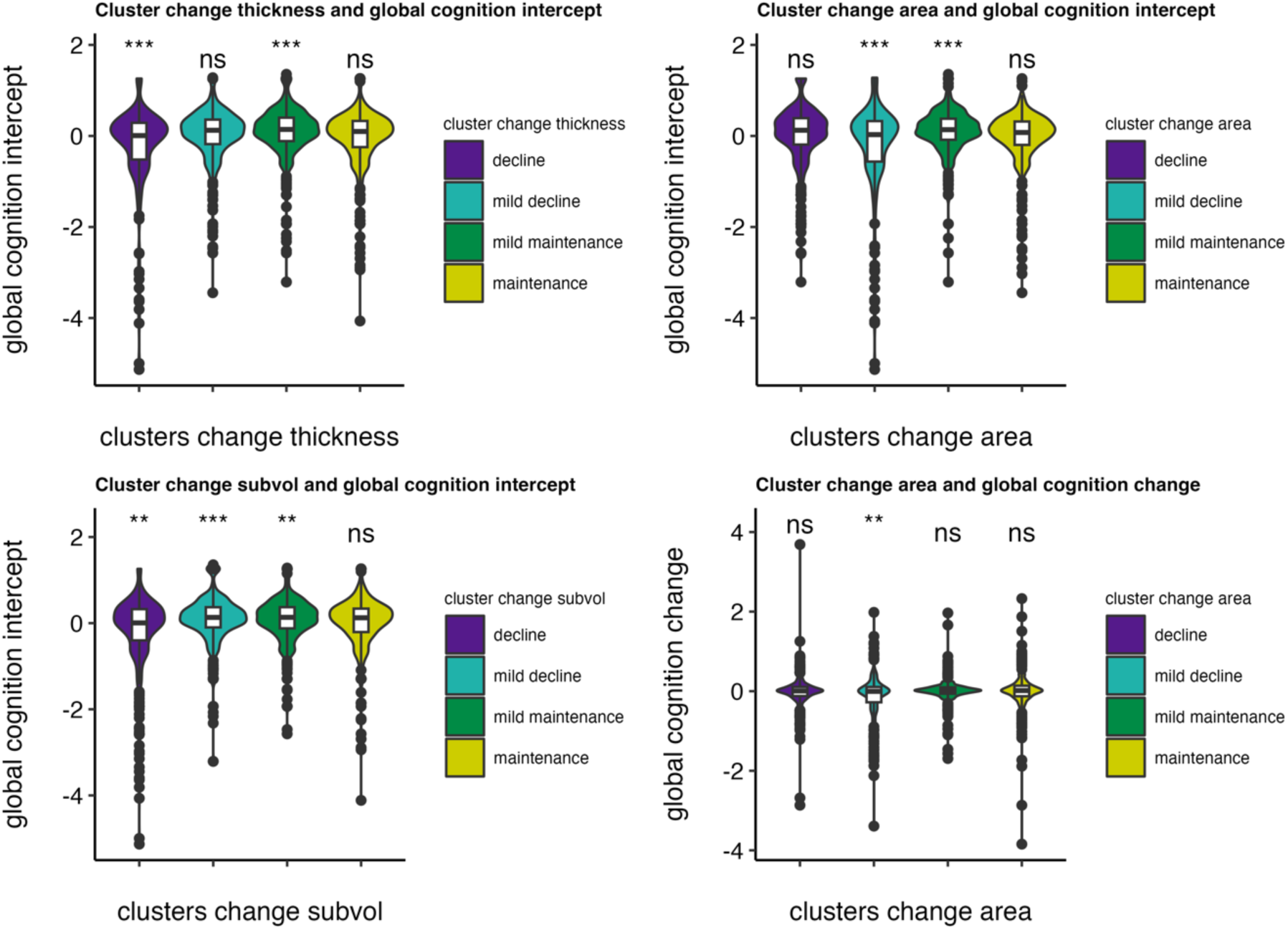
Significant associations between brain cluster assignment and global cognition. Post-hoc comparisons of means (one cluster vs. all) for global cognition intercept and change. ** = p < 0.01, *** = p < 0.001, ns = non-significant. Note that the term "global cognition" specifically pertains to the aforementioned screening tests.

**Table 3.** Associations between brain cluster assignment and cognitive functions.

| | Cognitive function | (F [pFDR]) | $\eta^2$ partial |
| --- | --- | --- | --- |
| Cluster change thickness | Memory change | 3.40 (0.10) | $7.05 \times 10^{-3}$ |
| | Memory intercept | 0.77 (0.67) | $1.61 \times 10^{-3}$ |
| | Global cognition change | 1.73 (0.16) | $3.35 \times 10^{-3}$ |
| | Global cognition intercept | 16.21 ($< 0.001$)<sup>(-a,c)</sup> | 0.02 |
| **Cluster change area** | Memory change | 0.85 (0.67) | $1.78 \times 10^{-3}$ |
| | Memory intercept | 2.64 (0.14) | $5.50 \times 10^{-3}$ |
| | Global cognition change | 4.16 (0.009)<sup>(-b)</sup> | $7.71 \times 10^{-3}$ |
| | Global cognition intercept | 15.69 ($< 0.001$)<sup>(-b,c)</sup> | 0.02 |
| **Cluster change subvolume** | Memory change | 0.51 (0.67) | $1.08 \times 10^{-3}$ |
| | Memory intercept | 0.52 (0.67) | $1.09 \times 10^{-3}$ |
| | Global cognition change | 1.84 (0.16) | $3.62 \times 10^{-3}$ |
| | Global cognition intercept | 15.88 ($< 0.001$)<sup>(-a,b,c)</sup> | 0.02 |

ANOVA models on the LME models output. Sex and Age at baseline (mean-centered) as covariates of no interest. Statistics represent F-values, pFDR corrected values, and $\eta^2$ partial represents the effect size eta squared partial. The superscripts represent the significant output ( $p < 0.01$ ) of the post hoc multiple comparisons, where each cluster assignment is compared against all: a) decline cluster $>$ mean; b) mild decline $>$ mean; c) mild maintenance $>$ mean; d) maintenance $>$ mean. A negative sign indicates that the significant comparison was lower than the mean. Please note that the term "global cognition" specifically pertains to the aforementioned screening tests.

### 3.5 Associations between brain cluster assignment and CSF AD biomarkers

The results are presented in **Table 4** (n = 612). Changes in subcortical volume clusters were significantly related to CSF Aβ42 (F = 8.30, df1= 605, df2 = 3, pFDR < 0.001), p-tau (F = 3.95, df1 = 600, df2 = 3, pFDR = 0.01), and p-tau/Aβ42 ratio (F = 10.40, df1 = 601, df2 = 3, pFDR < 0.001). We also found significant positive associations between changes in thickness clusters and the p-tau/Aβ42 ratio (F = 6.44, df1 = 601, df2 = 3, pFDR < 0.001) and Aβ42 (F = 7.08, df1 = 605, df2 = 3, pFDR < 0.001). Changes in cortical area clusters were significantly related to CSF Aβ42 (F = 4.54, df1 = 605, df2 = 3, pFDR = 0.007). The ANOVA results were in the expected direction, with clusters displaying more brain decline showing lower Aβ42, higher p-tau, and higher p-tau/Aβ42 ratio, and those showing relative brain maintenance exhibiting lower p-tau/Aβ42 ratio, and higher Aβ42. See **Supplementary Table 3**, **Figure 7** for the association with CSF AD biomarkers, and **Table 4** and **Supplementary Table 2** for the post-hoc comparisons across clusters, showing which specific subgroups had significantly different values.

**Figure 7.**
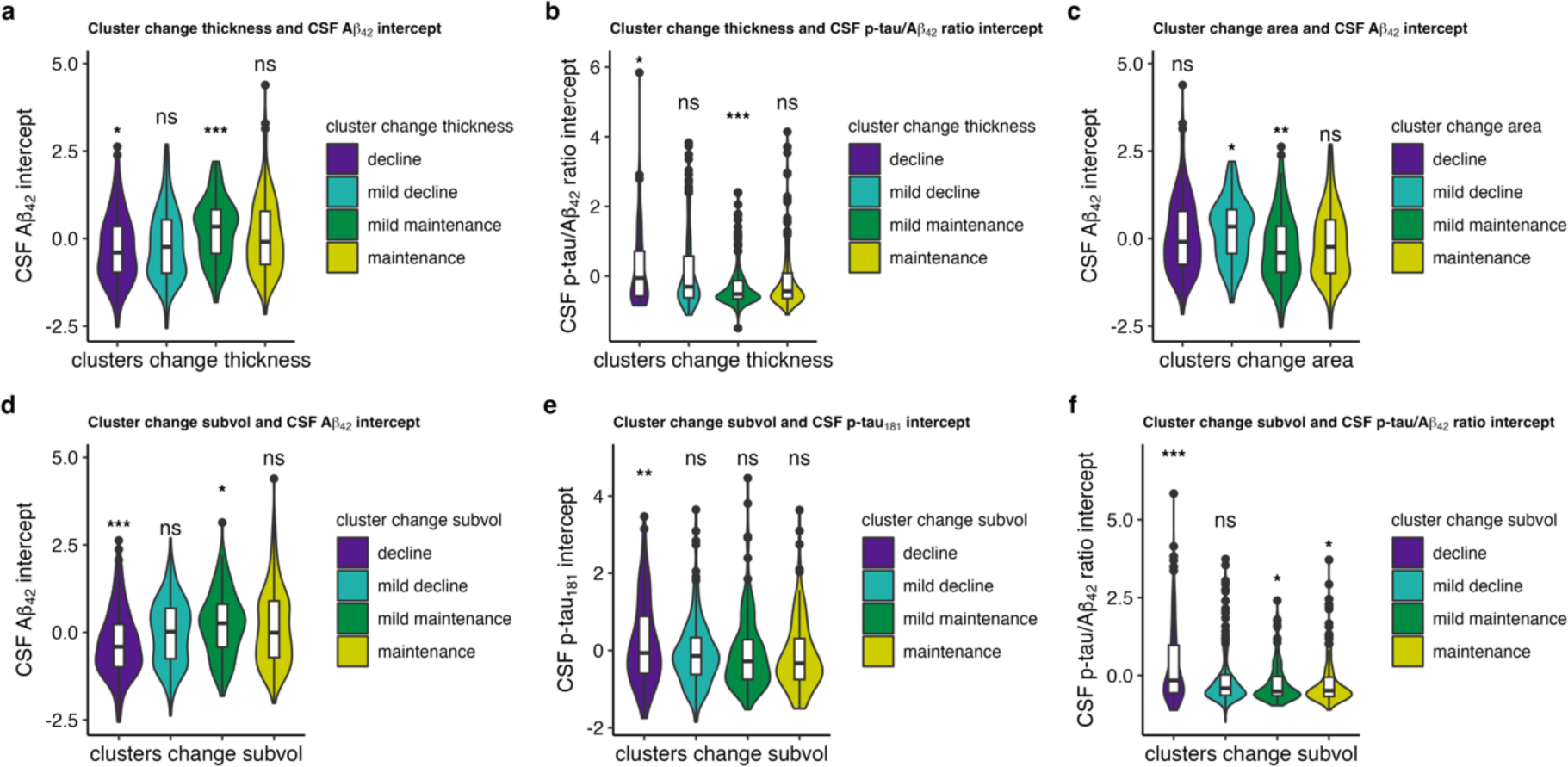
Significant associations between brain cluster assignment and AD CSF biomarkers. Post-hoc comparisons of means (one cluster vs all) for AD CSF biomarkers intercept. * = p < 0.05, ** = p < 0.01, *** = p < 0.001, ns = non-significant.

**Table 4.**
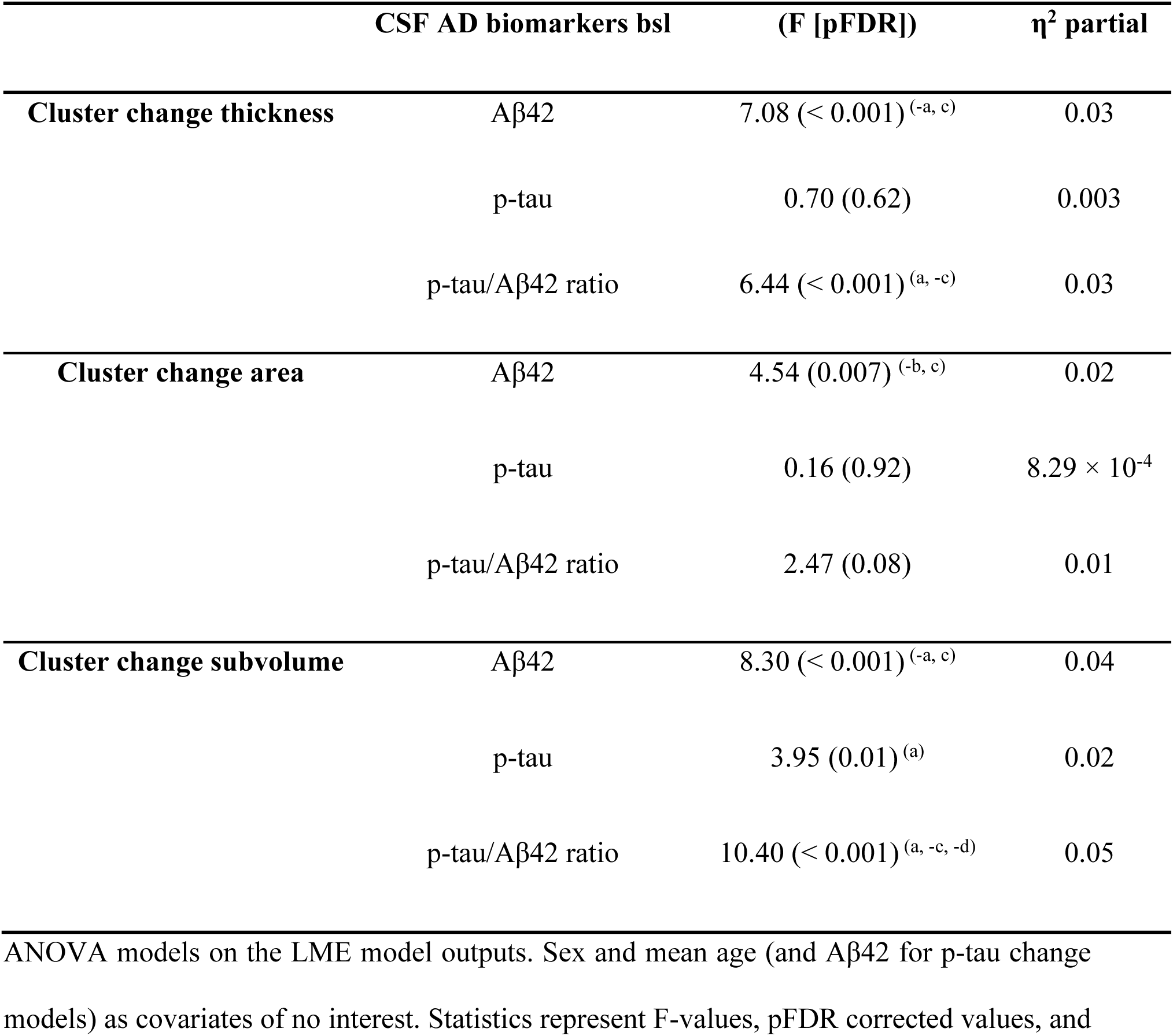
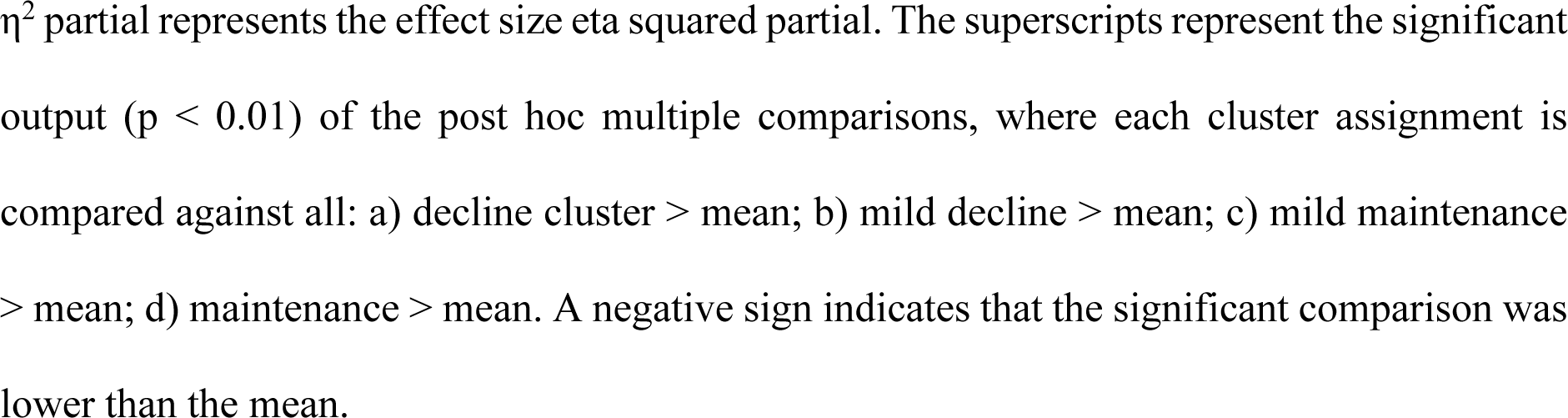
Associations between brain cluster assignment and CSF AD biomarkers at baseline.

### 3.6 Automated model selection for cognitive functioning and CSF core AD biomarkers

When using λ within 1 standard error of the minimum (the more conservative criterion), the LASSO models dropped all the predictors. By selecting the less conservative criteria for selecting λ (λ.min = 0.01), the optimal model for predicting the memory intercept included only the cluster area change, which had nonzero coefficients. The best model predicting memory change included the main effects of changes in thickness and subcortical volume, and the interactions among the three brain features (λ.min = 0.003). The optimal model predicting global cognition at baseline comprised thickness, area, and subcortical volume change main effects (λ.min = 0.004), whereas global cognitive change included area and subcortical volume main effects and their interactions (λ.min = 0.002). Regarding CSF AD biomarkers, we found that the main effects of changes in thickness, area, and subcortical volume were associated with CSF Aβ42 at baseline (λ.min = 0.01). Changes in subcortical volume best predicted p-tau (λ.min = 0.01), whereas the main effects of changes in thickness and subcortical volume were associated with the p-tau/Aβ42 ratio (λ.min = 0.01). See **Supplementary Table 4** for all the stats.

## 4. Discussion

We identified four ageotypes for cortical thickness, cortical area, and subcortical volume, grouping participants based on the degree of morphometric change. The overlap across modalities was low, indicating that a comprehensive understanding of structural brain changes in aging requires the integration of different brain features. The analysis of the associations between brain changes and cognitive function, as well as AD biomarkers, was beneficial in comprehending the significance of these brain changes in normal aging. In particular, clustering based on subcortical volumetric change was found to be highly sensitive to both cognition and AD biomarkers. This suggests that ageotypes are relevant in understanding cognitive decline in aging. Furthermore, the relationship with AD biomarkers indicates that structural brain changes may give rise to an increased risk for later development of AD.

Clustering was strongly based on a main factor of decline, suggesting that differences in cluster assignment could be attributed to a main “global” component of (modality-specific) brain decline rather than to specific spatial patterns. This finding is consistent with a previous study that used factor analysis on longitudinal volumetric ROIs changes and identified a general factor of cortical volume change in aging (Cox et al., 2021) which accounted for 63% of the longitudinal changes in the different regions. Similarly, Sele and colleagues (2020) found that a component of decline (from PCA) accounted for approximately 35% of the longitudinal volumetric change (slope differences) across different regions, especially temporal.

Although clusters were primarily determined by a global component of brain decline, some regions were especially critical for cluster assignment. Specifically, we found that subtypes based on both cortical thickness and cortical area change were strongly related to the degree of bilateral decline in the temporal and inferior parietal regions. These regions are among those suffering steeper age-related decline (Fjell et al., 2014b; Thambisetty et al., 2010), as well as exhibiting higher inter-individual variability (Sele et al., 2020). Notable decline in these regions can be seen also independently of *APOE* status and neurodegenerative processes reflected by AD biomarkers Aβ42 and tau, and is often considered characteristic of normal aging trajectories (Fjell et al., 2014a). Despite being highly vulnerable to aging, frontal regions did not have a special influence in determining cluster assignment, with the exception of the superior frontal cortex in cortical area change. One possible explanation is that despite showing a steep decline, these regions also showed relatively low inter-individual variability in change.

In other words, older participants tended to show a similar degree of change in these regions (Sele et al., 2021). Finally, we found that inter-individual variability in bilateral hippocampal volume decline and enlargement of the lateral ventricles was relevant for identifying clusters of subcortical volume changes. These regions are both strongly affected by age (Fjell et al., 2014a; Takao et al., 2012) with a high degree of variability across individuals (Sele et al., 2021, 2020) and are also commonly affected by AD (Apostolova et al., 2012; Grundman et al., 2002; Thompson et al., 2004).

Although individuals can be differentiated based on the main component of change within modality, the different modalities provide largely independent information in the context of age-related changes. A comprehensive approach that incorporates multiple measures of brain morphometric changes is essential to understand structural brain changes in older age. Indeed, there was minimal overlap in terms of cluster assignment among the brain features, particularly for cortical thickness and area. These two measures of surface, which together define cortical volume, are among other things thought to reflect the total number of cortical columns (area) and the number of cells within a column (thickness) (Rakic, 1988), respectively. Both area and thickness change are affected by increasing age, as shown by a cross-sectional and longitudinal study (Hogstrom et al., 2013; Storsve et al., 2014), and show a constant negative relationship across the adult lifespan (Storsve et al., 2014). Furthermore, these measures have distinct contributions to the volumetric changes at different stages of life. During development, cortical area changes play a significant role, and cortical thinning is the primary contributor in older age (Walhovd et al., 2016). Nevertheless, they showed an opposite pattern within regions; that is, those regions characterized by more thinning showed less decrease in area, and vice versa. Sele and colleagues (2021) found both null and negative associations between cortical area and cortical thickness change across individuals. Overall, these brain features show a unique genetic signature (Panizzon et al., 2009), although recently other researchers have reported opposing effects on the impact of genetics on thickness and area (Grasby et al., 2020), and might display specific biological processes that may account for the varying contributions to age-related structural changes. Therefore, although individuals can be differentiated based on the main component of change within modality, the different modalities provide largely independent biological information in the context of age-related changes.

Our findings showed that participants displaying more thinning, more subcortical volume decline, and/or more cortical area loss showed worse global cognition at baseline. These results can be interpreted in two ways. First, integrating brain reserve (Katzman et al., 1988; Stern et al., 2019) and maintenance (Nyberg et al., 2012) frameworks together within the Matthew principle. The latter posits an interaction between variation in level and change to explain differences in brain and cognition; in other words, it suggests that individuals who begin with an advantage will accumulate and maintain more advantage over time, and vice versa. From this perspective, participants with higher cognition at baseline may have accumulated neural resources that allowed them to counterbalance the effect of age-related brain changes. Consequently, the more neural resources available at our starting point (brain reserve), which accumulate over time, the more the advantages over time, leading to maintenance of brain resources available in aging, which is translated into better cognitive performance in older age. However, education, one of the most popular proxies of cognitive reserve (Stern, 2012) used to explain individual differences in cognition, which correlates with higher cognition in aging, does not seem to have a meaningful impact on structural brain changes in aging (Nyberg et al., 2021), and does not affect the relationship between brain change and cognitive change (Lovden et al., 2023), as would be predicted from the cognitive reserve account. Another alternative interpretation is that the relationship between global cognition and brain changes may capture the ongoing changes in the brain and cognition that occur prior to, during, and maybe even after the follow-up period. In other words, the follow-up period can be viewed as a temporal "window" for observing slow trajectories of the brain and cognitive decline. Indeed, we found change in cortical surface area is related to both baseline cognition and cognitive change (as assessed by screening parameters). Further, even the screening tests used assess global cognition, they cannot be considered a pre-morbid cognitive assessment. Thus, brain change –baseline cognition relationships seem to reflect a dynamic sluggish association of paired cognitive and brain change. This might indicate that the global cognition factor captures changes that occurred prior to neuroimaging acquisition and cannot be accounted for by earlier factors. A recent paper (Walhovd et al., 2023) argues that the timing of lifespan influences is crucial to explain individual differences in brain and cognition. In fact, it appears that differences in the trajectories of change in brain and cognition can only partially explain the inter-individual variability in older age. Instead, individual differences may be largely attributed to early life factors that remain relatively stable over the adult lifespan.

Cortical area changes were significantly related to cognitive changes in contrast to cortical thickness. Cortical area typically may indicate the number of cortical columns and it is related to information-processing capacity, and this was observed in older adults who showed cortical area decline, as they also exhibited more decline in the global cognition factor over time. This finding is supported by other studies (Borgeest et al., 2021; Nyberg et al., 2023), although they used fluid cognition measures (assessed by a speed of processing test and Cattell Culture Fair test). Changes in cortical thickness may be likely due to dendritic atrophy, which occurs with increasing age, and late-onset lower cortical thickness is associated with cognitive decline (de Chastelaine et al., 2019). We speculate that we did not find any positive association between thinning and cognitive change within our temporal interval due to the inclusion of relatively *young* older adults (aged 50 years and older). This may lead to relatively minor changes in cortical thickness, which accelerate with higher age, especially after 60 years, as shown in a previous study (Nyberg et al., 2023), where the association with cognitive change was significant only at the final time point, when participants were older. As we can see, the time interval is a critical factor in this context, and it is possible that both brain and cognitive changes occur simultaneously in the same time frame, or, as we speculate in our case, cognitive changes occur both prior to and later than our follow-up period. The global cognition factor, as measured in our case by the MMSE and MOCA scores, appears to be an earlier and valid predictor, capturing more general and systematic changes in the aging-disease continuum compared to memory alone, which generally encompasses more specific and subtle changes. Indeed, we did not observe any effect on memory. The relationship between MTL thinning and hippocampal volume decline with memory changes is well established (Fjell et al., 2014b; Gorbach et al., 2017; Leong et al., 2017). Hence, a possible explanation for this null association might be due to the memory – brain associations being more regionally specific (e.g., medial temporal lobe) than global cognitive scores.

Our results showed that more rapid cortical thinning, subcortical volume, and cortical area decline over time were related to lower CSF Aβ42 levels at baseline. Previous studies have reported conflicting results regarding the association between CSF Aβ42 and brain atrophy in cognitively healthy older adults (Fjell et al., 2014a; Svenningsson et al., 2019; Tosun et al., 2011; Wang et al., 2015). Indeed, some studies found that decreased Aβ42 levels were associated with hippocampal loss but not cortical thinning in AD-signature regions (Pettigrew et al., 2016; Wang et al., 2015). Conversely, another study (Arenaza-Urquijo et al., 2013) found cortical thinning in AD-vulnerable regions, while another cross-sectional study found no relationship between CSF Aβ42 positivity, hippocampal volume decline, or cortical thickness (Svenningsson et al., 2019). A significant association between lower CSF Aβ42 and surface area decline has not previously been reported. In our study, change in each longitudinal brain feature was associated with Aβ42. In addition, subcortical volumetric change was associated also to p-tau. Specifically, participants in the subcortical decline cluster, who showed higher p-tau and p-tau/Aβ42 ratio, as well as lower Aβ42 levels, may be at an increased risk for a subsequent clinical diagnosis of AD. Therefore, the association with AD biomarkers helps us understand the significance of these structural brain changes in the context of normal aging. Changes in the hippocampal volume and lateral ventricles are affected early in the disease process as long as AD biomarkers accumulate in the brain (Stricker et al., 2012). Overall, clustering of subcortical volume changes may provide helpful information for identifying individuals with an increased risk for a later clinical AD diagnosis, whereas the clustering of cortical features such as thickness and area may reflect different age-related brain processes.

### 4.1 Limitations and technical considerations

A strength of the present study is the use of longitudinal data for structural MRI, cognitive assessment, and CSF, which allows for a better capture of intra-individual changes over time. However, longitudinal studies can be affected by selective attrition, which means that results apply to the participants who did not drop out of the studies, who are known to be healthier, more educated, and with higher general cognitive ability than the general population (Beller et al., 2022; Salthouse, 2014). An additional problem with longitudinal data is the less than perfect reliability of the brain and cognitive change estimates. This aspect may help explain the stronger associations between brain change and baseline cognition compared to cognitive change. Additionally, it can be speculated that there is less variation in change than in level, making it more challenging to detect any systematic relationship. Another critical methodological aspect of this study is the merging of multiple cohorts, yielding increased statistical power and reduced sampling bias compared to meta-analytical approaches. However, this approach may also introduce new sources of error due to differences in measurements or populations (Zuo et al., 2019). This decision leads to the use of different memory and global cognitive tests across the different cohorts, and may lead to small biases because the same underlying construct is not necessarily captured.

## 5. Conclusions

In summary, this study identified four distinct ageotypes based on the global pattern of brain changes within cortical thickness, cortical area and subcortical volume measures over time. The minimal overlap across modalities highlights the need to combine all the features to better capture and understand age-related brain changes. Furthermore, the clustering of regional brain changes proved to be a valuable tool for explaining cognitive and biomarker differences in cognitively unimpaired older adults.

## Funding and acknowledgements

This work was supported by the Department of Psychology, University of Oslo (to K.B.W., A.M.F.), the Norwegian Research Council (to K.B.W., A.M.F., D.V.P [ES694407]) and the project has received funding from the European Research Council’s Starting Grant scheme under grant agreements 283634, 725025 (to A.M.F.). HZ is a Wallenberg Scholar and a Distinguished Professor at the Swedish Research Council supported by grants from the Swedish Research Council (#2023-00356; #2022-01018 and #2019-02397), the European Union’s Horizon Europe research and innovation programme under grant agreement No 101053962, Swedish State Support for Clinical Research (#ALFGBG-71320), the Alzheimer Drug Discovery Foundation (ADDF), USA (#201809-2016862), the AD Strategic Fund and the Alzheimer’s Association (#ADSF-21-831376-C, #ADSF-21-831381-C, #ADSF-21-831377-C, and #ADSF-24-1284328-C), the Bluefield Project, Cure Alzheimer’s Fund, the Olav Thon Foundation, the Erling-Persson Family Foundation, Stiftelsen för Gamla Tjänarinnor, Hjärnfonden, Sweden (#FO2022-0270), the European Union’s Horizon 2020 research and innovation programme under the Marie Skłodowska-Curie grant agreement No 860197 (MIRIADE), the European Union Joint Programme – Neurodegenerative Disease Research (JPND2021-00694), the National Institute for Health and Care Research University College London Hospitals Biomedical Research Centre, and the UK Dementia Research Institute at UCL (UKDRI-1003). KB is supported by the Swedish Research Council (2017-00915 and 2022-00732), the Alzheimer Drug Discovery Foundation (ADDF), USA (RDAPB-201809-2016615), the Swedish Alzheimer Foundation (AF-930351, AF-939721 and AF-968270), Hjärnfonden, Sweden (FO2017-0243 and ALZ2022-0006) the Swedish state under the agreement between the Swedish government and the County Councils, the ALF-agreement (ALFGBG-715986 and ALFGBG-965240), the European Union Joint Program for Neurodegenerative Disorders (JPND2019-466-236), the National Institute of Health (NIH), USA, (grant 1R01AG068398-01), and the Alzheimer’s Association 2021 Zenith Award (ZEN-21-848495) and the Alzheimer’s Association 2022-2025 Grant (SG-23-1038904 QC). L.O.W. and data collection in COGNORM is funded by the South-Eastern Norway Regional Health Authorities (#2017095) The Norwegian Health Association (#19536) and by Wellcome Leap’s Dynamic Resilience Program (jointly funded by Temasek Trust) #104617). The funding sources had no role in the study design. Data collection and sharing for this project were funded by the ADNI (NIH Grant U01 AG024904) and DOD ADNI (Department of Defense award number W81XWH-12-2-0012). ADNI is funded by the National Institute on Aging, the National Institute of Biomedical Imaging and Bioengineering, and through generous contributions from the following: AbbVie, Alzheimer’s Association; Alzheimer’s Drug Discovery Foundation; Araclon Biotech; BioClinica, Inc.; Biogen; Bristol-Myers Squibb Company; CereSpir, Inc.; Cogstate Eisai Inc.; Elan Pharmaceuticals, Inc.; Eli Lilly and Company; EuroImmun; F. Hoffmann-La Roche Ltd and its affiliated company Genentech, Inc.; Fujirebio; GE Healthcare; IXICO Ltd.; Janssen Alzheimer Immunotherapy Research & Development, LLC.; Johnson & Johnson Pharmaceutical Research & Development LLC.; Lumosity; Lundbeck; Merck & Co., Inc.; Meso Scale Diagnostics, LLC.; NeuroRx Research; Neurotrack Technologies; Novartis Pharmaceuticals Corporation; Pfizer Inc.; Piramal Imaging; Servier; Takeda Pharmaceutical Company; and Transition Therapeutics. The Canadian Institutes of Health Research is providing funds to support ADNI clinical sites in Canada. Private sector contributions are facilitated by the Foundation for the National Institutes of Health (http://www.fnih.org). The grantee organization is the Northern California Institute for Research and Education, and the study is coordinated by the Alzheimer’s Therapeutic Research Institute at the University of Southern California. ADNI data are disseminated by the Laboratory for Neuro Imaging at the University of Southern California. The investigators within the ADNI contributed to the design and implementation of ADNI and/or provided data but did not participate in analysis or writing of this report. A complete listing of ADNI investigators can be found at: http://adni.loni.usc.edu/wp-content/uploads/how_to_apply/ADNI_Acknowledgement_List.pdf. OASIS 3 data were provided [in part] by OASIS 3: Longitudinal Multimodal Neuroimaging: Principal Investigators: T. Benzinger, D. Marcus, J. Morris; NIH P30 AG066444, P50 AG00561, P30 NS09857781, P01 AG026276, P01 AG003991, R01 AG043434, UL1 TR000448, R01 EB009352. AV-45 doses were provided by Avid Radiopharmaceuticals, a wholly owned subsidiary of Eli Lilly. Some of the data used in preparation of this article were obtained from the PREVENT-AD program (Breitner et al., 2016). PREVENT-AD was funded by the Canadian Institutes of Health Research, McGill University, the Fonds de Recherche du Québec – Santé, Alzheimer’s Association, Brain Canada, the Government of Canada, the Canada Fund for Innovation, the Douglas Hospital Research Centre and Foundation, the Levesque Foundation, an unrestricted research grant from Pfizer Canada. Some of the data used in the preparation of this article was obtained from the AIBL. AIBL is funded by the Commonwealth Scientific and Industrial Research Organisation (CSIRO), which was made available at the ADNI database (www.loni.usc.edu/ADNI). The AIBL researchers contributed data but did not participate in analysis or writing of this report. AIBL researchers are listed at www.aibl.csiro.au. Some of the data used in the preparation of this article were obtained from the Harvard Aging Brain Study (HABS - P01AG036694; https://habs.mgh.harvard.edu). HABS data release 2.0, obtained August, 2022 via habs.mgh.harvard.edu. The HABS study was launched in 2010, funded by the National Institute on Aging, and is led by principal investigators Reisa A. Sperling MD and Keith A. Johnson MD at Massachusetts General Hospital/Harvard Medical School in Boston, MA.

