## Supplementary Material for "Subtypes of brain change in aging and their associations with cognition and Alzheimer’s disease biomarkers"

**Participants**

<u>COGNORM</u>

The COGNORM cohort was recruited at the Oslo University Hospital and Diakonhjemmet Hospital, Oslo, Norway. Patients (age ≥ 65 years) scheduled for elective gynecological (genital prolapse), urological (benign prostate hyperplasia, prostate cancer, or bladder tumor/cancer), or orthopedic (knee or hip replacement) surgery under spinal anesthesia were recruited. Participants were required to have no dementia, previous stroke with sequela, Parkinson's disease, or other neurodegenerative diseases that are likely to affect cognition. CSF samples were collected by an anesthesiologist in conjunction with spinal anesthesia, and patients underwent magnetic resonance imaging (MRI) after surgery (interval = 60 days). Patients with suspected undiagnosed dementia at any time within the first five years of follow-up (n = 15) (Sajjad et al., 2020), MMSE score <28 at baseline, and at least two abnormal cognitive test scores (-1.5 standard deviation [SD] below the mean normal value for age, sex, and education) were excluded. All observations corresponding to cognitively normal individuals were included. Only individuals with two or more MRI observations were included in the sample. The final sample included 95 participants with available longitudinal MRI data.

<u>Alzheimer's Disease Neuroimaging Initiative (ADNI)</u>

ADNI is a multi-site project led by Doctor Michael W. Weiner to assess the progression of mild cognitive impairment (MCI) and early Alzheimer's Disease (AD), combining imaging, clinical and other biological markers, and neuropsychological and clinical assessments over time. The ADNI was launched in 2023 as a public-private partnership. For more information, visit https://adni.loni.usc.edu/about/. The age range for the participants is 55-90 years. In the present study, we included participants from ADNI 1 to ADNI 3, who were cognitively healthy at baseline, as evaluated by the ADNI team. Only observations in which participants were still cognitively healthy were included. Participants were required to have no evidence of ischemic stroke (Hachinski Ischemic Score ≤ 4), a Geriatric Depression scale score < 6, stable medications for 4 weeks before the screening, good auditory and visual acuity, good general health, no medical contraindications to MRI and at least 6 grades of education/work history. General inclusion and exclusion criteria are described elsewhere (Petersen et al., 2010). All participants signed an informed consent form. The protocols were approved by the corresponding ethical committees. The final sample included 544 participants with longitudinal MRI.

Open Access Series of Imaging Studies (OASIS 3)

The Open Access Series of Imaging Studies (OASIS) is a collection of multimodal data that focuses on the impact of healthy aging and AD and is openly accessible to the scientific community. OASIS-3, a specific subset of this collection, encompasses MRI and PET imaging data, along with related clinical data, for 1098 participants at the Washington University Knight Alzheimer Disease Research Center. Participants included 605 cognitively normal adults and 493 individuals with various stages of cognitive decline. The age range of the patients was 42–95 years. See the general inclusion and exclusion criteria (LaMontagne et al., 2019). All participants consented to Knight ADRC-related projects following procedures approved by the Institutional Review Board of Washington University School of Medicine. Participants aged < 65 years underwent clinical and cognitive assessments every 3 years, while participants aged ≥ 65 years underwent annual clinical and cognitive assessments. Only participants deemed cognitively normal at baseline were included in the observations. Observations were included until the last observation in which a subject was deemed cognitively healthy. The final sample comprised 518 participants with longitudinal MRI.

The Australian Imaging, Biomarker and Lifestyle Study (AIBL)

The AIBL Study is a prospective study of 1,112 individuals, including 768 cognitively normal participants, 133 with mild cognitive impairment, and 211 with AD. These participants undergo detailed assessments every 18 months, and an additional 1,247 individuals have been added as an "enrichment cohort" over the past decade. The age range of the participants was 60–96 years old. The study focuses on AD biomarkers, utilizing techniques such as MRI, PET, blood tests, and CSF analysis, as well as neuropsychological and clinical assessments. Healthy participants must meet specific criteria, including being free of cognitive impairments and having test performance within 1.5 SD of age-adjusted norms. The test battery and sample description have been described in detail previously (Ellis et al., 2009). In the present study, the AIBL team defined the clinical criteria. All observations corresponding to cognitively normal individuals were included. Only individuals with two or more MRI observations were included in the sample. Our final sample included 149 participants with longitudinal MRI information.

Harvard Aging Brain Study (HABS)

HABS is a long-term observational study that aims to enhance our understanding of brain aging and the early stages of Alzheimer's disease. The study involves the use of PET, MRI data (collected at 3 and 5 years), as well as annual neuropsychological and clinical assessments. The study comprised 290 participants who were followed up for up to five years after the baseline assessment. The age range was 62-90 years at the time of baseline assessment, and all patients were considered non-clinically impaired at the start of the study. To be included in the study, participants were $\geq$ 65 years old, had a CDR score of 0, MMSE score $\geq$ 25, < 11 on the Geriatric

Depression Scale, and scores above age- and education-adjusted cutoffs on the 30-Minute Delayed Recall of the Logical Memory Story A. Participants with a history of alcoholism, drug abuse, head trauma, or current serious medical/psychiatric illness were excluded. Further details can be found elsewhere (Dagley et al., 2017). In our study, we included observations in which participants were considered non-clinically impaired. Only individuals with 2 or more observations with MRI data were included. The final sample study comprised 166 participants with longitudinal MRI.

Pre-symptomatic Evaluation of Experimental or Novel Treatments for AD (PREVENT-AD)

The PREVENT-AD is a long-term study that follows cognitively healthy older individuals with a familiar history of AD. It includes 348 participants, either from the observational cohort or from the main clinical trial of PREVENT-AD. This study comprises MRI images, blood and CSF samples, and clinical and neuropsychological assessments. Participants in the study had to be at least 60 years old (with the exception of individuals between 55 and 59 years old, who were eligible if their age was within 15 years of symptom onset of their youngest affected first-degree relative). These participants had to have at least 6 years of education, and they needed to be cognitively unimpaired at baseline. The Montreal Cognitive Assessment (MoCA) and CDR scales were used to assess cognitive abilities, and participants were considered cognitively intact if their MoCA scores were $\geq$ 26/30 or their CDR was = 0. If they were cognitively intact at baseline (and a complete neuropsychological assessment was performed if there was any doubt), the Repeatable Battery for Assessment of Neuropsychological Status (RBANS) was administered annually. If their cognitive performance was lower than expected or their RBANS index score was >1 SD below the mean in the two different cognitive domains at any visit, they were referred for neuropsychological evaluation. If this evaluation suggested probable mild cognitive impairment (MCI), they were excluded from the study. Other exclusion criteria at baseline included medical conditions that prevented longitudinal participation or medical contraindications to MRI, use of acetylcholinesterase inhibitors, other approved prescription cognitive enhancers, hypertension, or substance abuse. The inclusion and exclusion criteria have been previously described in detail (Tremblay-Mercier et al., 2021). The protocols, consent forms, and study procedures were approved by the McGill Institutional Review Board and/or Douglas Mental Health University Institute Research Ethics Board. In our study, we included observations in which participants were considered non-clinically impaired. Only individuals with 2 or more observations with MRI data were included. The final sample consisted of 229 participants.

Center for Lifespan Changes in Brain and Cognition (LCBC)

The LCBC study involved recruiting cognitively healthy individuals from the community across different age ranges through a variety of methods such as newspapers and webpage ads. All adult participants had to undergo a standardized health interview before being included in the study, and those with a history of neurological or psychiatric conditions or who reported concerns about their cognitive function were excluded. Additionally, all participants over the age of 40 years were required to score at least 24 on the Mini-Mental State Examination. Further details can be found elsewhere (Fjell et al., 2023). In our study, we included observations in which participants were considered non-clinically impaired. Only individuals with 2 or more observations with MRI data were included. The final sample study comprised 198 participants with longitudinal MRI.

**Supplementary Table MRI acquisition parameters**

| Sample | Scanner | Field strength (Tesla) | Sequence parameters |
|---|---|---|---|
| COGNORM | Siemens Avanto scanner *12-channel coil* | 1.5 | TR: 2400 ms, TE: 3.79 ms, TI: 1000 ms, flip angle: 8°, voxel size: 1.25 × 1.25 × 1.20 mm$^3$, FoV: 240 mm |
| ADNI (1-2-3-GO) | Variable | 1.5 and 3.0 | See https://adni.loni.usc.edu/methods/documents/mri-protocols/ |
| OASIS 3 | Siemens Vision, TIM Trio, Sonata, and BioGraph mMR *16-channel coil* | 1.5 and 3.0 | TR: 9.7 ms, TE: 4.0 ms, TI: 20 ms, flip angle: 10°, FoV: 256×256 mm<br><br>TR: 2400 ms, TE: 3.1 ms, TI: 1000 ms, flip angle: 8°, voxel size: 1 × 1 × 1 mm$^3$, FoV: 256×256 mm<br><br>TR: 9.7 ms, TE: 3.9 ms, TI: 20 ms, flip angle: 15°, FoV: 224×256 mm<br><br>TR: 2300 ms, TE: 2.3 ms, TI: 900 ms, flip angle: 9°, FoV: 240×256 mm |
| AIBL | Siemens Avanto, Tim Trio and Verio | 1.5 and 3.0 | TR: 1900 ms /2300 ms, TE: 2.98 ms, TI: 900 ms, flip angle: 9°, voxel size: 1×1×1.2 mm$^3$, FoV: 240×256 mm |
| HABS | Siemens Trio-Tim *12-channel coil* | 3.0 | TR: 2300 ms, TE: 2.98 ms, TI: 900 ms, flip angle: 9°, voxel size: 1×1×1.2 mm$^3$, FoV: 240×256 mm<br><br>TR: 2200 ms, TE: 1.5/3.4/5.2/7.0 ms, TI: 1100 ms, flip angle: 7°, voxel size: 1×1×1.2 mm$^3$, FoV: 228×228 mm |
| PREVENT-AD | Siemens TIM Trio *12 or 32-channel coil* | 3.0 | TR: 2300 ms, TE: 2.98 ms, TI: 900 ms, flip angle: 9°, voxel size: 1×1×1 mm$^3$, FoV: 256×240×176 mm |
| LCBC | Siemens Prisma, Skyra, and Avanto | 1.5 and 3.0 | TR: 2400 ms, TE: 2.22 ms, TI: 1000 ms, flip angle: 8°, voxel size: 1×1×1, FoV: 240×256 mm (*Prisma*) |

TR: 2300 ms, TE: 2.98 ms, TI: 850 ms, flip angle: 8°, voxel size: 1×1×1, FoV: 256×256 mm *(Skyra)*

TR: 2400 ms, TE: 3.79 ms, TI: 1000 ms, flip angle = 8°, voxel size: 1.25×1.25×1.25, FoV: 240 × 240 mm *(Avanto)*

Specific information on image acquisition for the different samples. All the scans acquired are T1 weighted structural scans, MPRAGE scans. TR: Repetition time, TE: Echo time, TI: Inversion time, FoV: Field of View.

**Supplementary Table 1 Post-hoc comparisons for the significant associations between brain cluster assignment and genetic-environmental variables**

| | Thickness | Area | Subcortical volume |
|---|---|---|---|
| **Sex** | X | *Mild decline cluster* males over-represented (residuals = 2.43)<br>*Mild maintenance cluster* males under-represented (residuals = -3.82) | *Decline cluster* males over-represented (residuals = 4.38)<br>*Mild maintenance cluster* males under-represented (residuals = - 3.82) |
| **Education** | X | X | X |
| **Age** | *Decline cluster* > higher age (β = 2.07, p = 0.002)<br>*Mild maintenance cluster* < lower age (β = -1.89, p < 0.001) | *Mild decline cluster* > higher age (β = 3.12, p < 0.001)<br>*Mild maintenance cluster* < lower age (β = -3.25, p < 0.001) | *Decline cluster* > higher age (β = 1.88, p < 0.001)<br>*Mild decline cluster* < lower age (β = -1.77, p < 0.001)<br>*Mild maintenance cluster* < lower age (β = -1.62, p < 0.001)<br>*Maintenance cluster* > higher age (β = 1.55, p = 0.002) |
| ***APOE* ε4** | *Mild maintenance cluster APOE* ε4 under-represented (residuals = -2.01) | *Decline cluster APOE* ε4 carriers over-represented (residuals = 1.74)<br>*Maintenance cluster APOE* ε4 carriers under-represented (residuals = - 1.87) | *Decline cluster APOE* ε4 carriers over-represented (residuals = 3.01) |

| | | | |
|---|---|---|---|
| **Cohort** | *Decline cluster* over-represented ADNI (residuals = 3.85), under-represented LCBC (residuals = -3.18) and HABS (residuals = -2.39)<br>*Mild maintenance cluster* over-represented LCBC (residuals = 4.38), under-represented ADNI (residuals = -3.99)<br>*Maintenance cluster* over-represented ADNI (residuals = 3.09), under-represented LCBC (residuals = -3.12) | *Decline cluster* over-represented PREVENT-AD (residuals = 3.86)<br>*Mild decline cluster* over-represented ADNI (residuals = 2.54), under-represented HABS (residuals = -3.50) and LCBC (residuals = -2.21)<br>*Mild maintenance cluster* over-represented NBM (residuals = 2.35), under-represented ADNI (residuals = -4.15)<br>*Maintenance cluster* over-represented ADNI (residuals = 3.83), under-represented PREVENT-AD (residuals = -5.72) | *Decline cluster* more PREVENT-AD (p = 0.001), less LCBC (p = 0.003) over-represented ADNI (residuals = 2.73), under-represented NBM (residuals = -2.34) and LCBC (residuals = -2.95)<br>*Mild decline cluster* over-represented PREVENT-AD (residuals = 3.40) and LCBC (residuals = 5.07), under-represented ADNI (residuals = -2.88) and OASIS (residuals = -2.36)<br>*Mild maintenance cluster* over-represented HABS (residuals = 4.22) and NBM (residuals = 2.46), under-represented ADNI (residuals = -3.43)<br>*Maintenance cluster* more ADNI (p < 0.001), OASIS-3 (p = 0.02), less PREVENT-AD (p = 0.002), LCBC (p < 0.001), HABS (p = 0.009) over-represented ADNI (residuals = 3.98) and OASIS (residuals = 2.49), under-represented HABS (residuals = -2.86), PREVENT-AD (residuals = -3.16), and LCBC (residuals = -4.15) |

We assessed whether the cluster assignment (for brain feature) differed for sex, age, education, *APOE* ε4 status, and Cohort. Post-hoc multiple comparisons for age and education were applied only to significant LME and ANOVA outputs, performing each cluster assignment against all. The output of multiple comparisons of means displayed the adjusted p-values using the single-step method. For sex, *APOE* ε4 status, and Cohort, we applied *chisq.residuals* for the adjusted residuals, which provides info about how much the variable of interest is under/over-represented among clusters and is used as a post-hoc measure for chi-square tests.

**Supplementary Table 2 Post-hoc comparisons for the significant associations between brain cluster assignment and variables of interest**

| | Thickness | Area | Subcortical volume |
|---|---|---|---|
| **Global cognition intercept** | *Decline cluster* (β = -0.28, $p < 0.001$*)<br>*Mild decline cluster* (β = 0.08, p = 0.06)<br>*Mild maintenance cluster* (β = 0.19, $p < 0.001$*)<br>*Maintenance cluster* (β = 0.01, p = 0.98) | *Decline cluster* (β = 0.08, p = 0.08)<br>*Mild decline cluster* (β = -0.27, $p < 0.001$*)<br>*Mild maintenance cluster* (β = 0.18, $p < 0.001$*)<br>*Maintenance cluster* (β = 0.008, p = 0.99) | *Decline cluster* (β = -0.23, $p < 0.001$*)<br>*Mild decline cluster* (β = 0.12, p = 0.001*)<br>*Mild maintenance cluster* (β = 0.11, p = 0.006*)<br>*Maintenance cluster* (β = -0.004, p = 0.99) |
| **Global cognition change** | X | *Decline cluster* (β = 0.02, p = 0.86)<br>*Mild decline cluster* (β = -0.10, p = 0.002*)<br>*Mild maintenance cluster* (β = 0.04, p = 0.19)<br>*Maintenance cluster* (β = 0.04, p = 0.24) | X |
| **Aβ42** | *Decline cluster* (β = -0.31, p = 0.02*)<br>*Mild decline cluster* (β = -0.18, p = 0.17)<br>*Mild maintenance cluster* (β = 0.35, $p < 0.001$*)<br>*Maintenance cluster* (β = 0.15, p = 0.30) | *Decline cluster* (β = -0.04, p = 0.96)<br>*Mild decline cluster* (β = -0.32, p = 0.02*)<br>*Mild maintenance cluster* (β = 0.30, p = 0.005*)<br>*Maintenance cluster* (β = 0.06, p = 0.86) | *Decline cluster* (β = 0.28, $p < 0.001$*)<br>*Mild decline cluster* (β = -0.03, p = 0.99)<br>*Mild maintenance cluster* (β = 0.28, p = 0.02*)<br>*Maintenance cluster* (β = 0.18, p = 0.17) |
| **p-tau** | X | X | *Decline cluster* (β = 0.29, p = 0.005*)<br>*Mild decline cluster* (β = 0.29, p = 0.99)<br>*Mild maintenance cluster* (β = -0.07, p = 0.86)<br>*Maintenance cluster* (β = -0.19, p = 0.10) |

| | | | |
|---|---|---|---|
| **p-tau/Aβ42 ratio** | *Decline cluster* (β = 0.28, p = 0.04*)<br>*Mild decline cluster* (β = 0.17, p = 0.17)<br>*Mild maintenance cluster* (β = -0.32, p < 0.001*)<br>*Maintenance cluster* (β = -0.13, p = 0.38) | X | *Decline cluster* (β = 0.47, p < 0.001*)<br>*Mild decline cluster* (β = 0.02, p = 0.99)<br>*Mild maintenance cluster* (β = -0.26, p = 0.02*)<br>*Maintenance cluster* (β = -0.22, p = 0.04*) |

Post-hoc comparisons for cognitive variables and CSF AD biomarkers were applied only to significant LME and ANOVA outputs, performing each cluster assignment against all. The output of the multiple comparisons of means displayed adjusted p-values using the single-step method, as implemented in the *multcomp* R-package (Westfall, 2010). β = estimate, p = adjusted p-values.

**Supplementary Table 3 Associations between brain cluster assignment and CSF AD biomarkers changes**

| | CSF AD biomarkers bsl | (F [pFDR]) | $\eta^2$ partial |
|---|---|---|---|
| **Cluster change thickness** | Aβ42 | 1.66 (0.31) | 0.01 |
| | p-tau | 0.45 (0.81) | 0.004 |
| | p-tau/Aβ42 ratio | 3.05 (0.13) | 0.03 |
| **Cluster change area** | Aβ42 | 2.47 (0.28) | 0.02 |
| | p-tau | 0.91 (0.65) | 0.008 |
| | p-tau/Aβ42 ratio | 3.77 (0.10) | 0.03 |
| **Cluster change subvolume** | Aβ42 | 0.27 (0.85) | 0.002 |
| | p-tau | 0.62 (0.77) | 0.006 |
| | p-tau/Aβ42 ratio | 1.81 (0.31) | 0.02 |

Longitudinal values available for 328 (Aβ42), 327 (p-tau), and 326 (p-tau/Aβ42 ratio) participants. We scaled the CSF values based on the mean and SD at the first timepoint within cohort. To compute the slope used for the analysis, we ran a linear regression model for each participant, with age as the predictor (difference between age at the time of CSF collection and age at the baseline of the first MRI measurement) and CSF scaled as outcome. We used LME to compute the effect of cluster assignment on CSF AD biomarker intercepts and changes. Additionally, 4-group ANOVA models were run on the outputs of the LME models. The models were corrected for multiple comparisons using the false discovery rate and Benjamini-Hochberg correction (pFDR) (Benjamini and Hochberg, 1995). In this table, we report the ANOVA results for the LME model output. Sex and mean age (and Aβ42 for p-tau change models) as covariates of no interest. Statistics represent F-values, pFDR-corrected values, and $\eta^2$ partial represents the effect size eta squared partial. No significant relationship between cluster assignment and AD biomarker changes survived correction for multiple comparisons using pFDR ($p > .05$).

**Supplementary Table 4 Automated model selection for cognitive functioning and CSF core AD biomarkers**

| | Memory bsl β | Memory change β | Global cognition bsl β | Global cognition change β | Aβ42 bsl β | p-tau bsl β | p-tau/Aβ42 ratio bsl β |
|---|---|---|---|---|---|---|---|
| (Intercept) | 0.05 | 0.08 | 0.05 | -0.01 | 0.06 | 0.03 | -0.05 |
| cluster delta cth2 | 0 | -0.08 | 0.05 | 0 | 0.12 | 0 | -0.14 |
| cluster delta cth3 | 0 | -0.22 | -0.08 | 0 | -0.09 | 0 | 0.08 |
| cluster delta cth4 | 0 | -0.02 | 0.03 | 0 | -0.10 | 0 | 0.07 |
| cluster delta area2 | -0.05 | 0 | -0.09 | -0.04 | -0.09 | 0 | 0 |
| cluster delta area3 | $-8.42 \times 10^{-3}$ | 0 | 0.03 | $7.82 \times 10^{-3}$ | 0.08 | 0 | 0 |
| cluster delta area4 | 0.04 | 0 | -0.02 | 0.02 | 0.02 | 0 | 0 |
| cluster delta vol2 | 0 | 0.04 | -0.11 | -0.01 | -0.21 | 0.12 | 0.26 |
| cluster delta vol3 | 0 | -0.01 | 0.03 | $6.25 \times 10^{-3}$ | -0.09 | -0.01 | 0.07 |
| cluster delta vol4 | 0 | 0.03 | $8.29 \times 10^{-3}$ | $6.33 \times 10^{-3}$ | 0.10 | -0.08 | -0.10 |
| Sex | -0.14 | -0.08 | -0.08 | $-6.08 \times 10^{-3}$ | $-6.19 \times 10^{-3}$ | -0.09 | -0.01 |
| Age Bsl | $-8.50 \times 10^{-3}$ | $5.68 \times 10^{-3}$ | $3.43 \times 10^{-3}$ | $-2.40 \times 10^{-3}$ | $-1.35 \times 10^{-3}$ | 0.03 | 0.02 |
| cluster delta cth2:cluster delta area2 | 0 | 0 | 0 | 0 | 0 | 0 | 0 |
| cluster delta cth3:cluster delta area2 | 0 | -0.02 | 0 | 0 | 0 | 0 | 0 |
| cluster delta cth4:cluster delta area2 | 0 | $-8.23 \times 10^{-3}$ | 0 | 0 | 0 | 0 | 0 |
| cluster delta cth2:cluster delta area3 | 0 | $-4.27 \times 10^{-3}$ | 0 | 0 | 0 | 0 | 0 |
| cluster delta cth3:cluster delta area3 | 0 | 0 | 0 | 0 | 0 | 0 | 0 |
| cluster delta cth4:cluster delta area3 | 0 | $2.19 \times 10^{-3}$ | 0 | 0 | 0 | 0 | 0 |

| | | | | | | | |
|---|---|---|---|---|---|---|---|
| cluster delta cth2:cluster delta area4 | 0 | -0.01 | 0 | 0 | 0 | 0 | 0 |
| cluster delta cth3:cluster delta area4 | 0 | $4.63 \times 10^{-3}$ | 0 | 0 | 0 | 0 | 0 |
| cluster delta cth4:cluster delta area4 | 0 | 0.02 | 0 | 0 | 0 | 0 | 0 |
| cluster delta cth2:cluster delta vol2 | 0 | -0.01 | 0 | 0 | 0 | 0 | 0 |
| cluster delta cth3:cluster delta vol2 | 0 | -0.04 | 0 | 0 | 0 | 0 | 0 |
| cluster delta cth4:cluster delta vol2 | 0 | $2.99 \times 10^{-3}$ | 0 | 0 | 0 | 0 | 0 |
| cluster delta cth2:cluster delta vol3 | 0 | 0.03 | 0 | 0 | 0 | 0 | 0 |
| cluster delta cth3:cluster delta vol3 | 0 | -0.02 | 0 | 0 | 0 | 0 | 0 |
| cluster delta cth4:cluster delta vol3 | 0 | -0.03 | 0 | 0 | 0 | 0 | 0 |
| cluster delta cth2:cluster delta vol4 | 0 | $-2.25 \times 10^{-3}$ | 0 | 0 | 0 | 0 | 0 |
| cluster delta cth3:cluster delta vol4 | 0 | 0.04 | 0 | 0 | 0 | 0 | 0 |
| cluster delta cth4:cluster delta vol4 | 0 | $-1.28 \times 10^{-4}$ | 0 | 0 | 0 | 0 | 0 |
| cluster delta area2:cluster delta vol2 | 0 | 0.04 | 0 | -0.03 | 0 | 0 | 0 |
| cluster delta area3:cluster delta vol2 | 0 | $-4.79 \times 10^{-3}$ | 0 | $6.72 \times 10^{-3}$ | 0 | 0 | 0 |
| cluster delta area4:cluster delta vol2 | 0 | 0.14 | 0 | 0.02 | 0 | 0 | 0 |
| cluster delta area2:cluster delta vol3 | 0 | -0.05 | 0 | $-2.28 \times 10^{-3}$ | 0 | 0 | 0 |
| cluster delta area3:cluster delta vol3 | 0 | -0.02 | 0 | 0.02 | 0 | 0 | 0 |
| cluster delta area4:cluster delta vol3 | 0 | 0.09 | 0 | $-8.84 \times 10^{-4}$ | 0 | 0 | 0 |
| cluster delta area2:cluster delta vol4 | 0 | -0.03 | 0 | $5.31 \times 10^{-3}$ | 0 | 0 | 0 |
| cluster delta area3:cluster delta vol4 | 0 | 0.04 | 0 | $-5.54 \times 10^{-5}$ | 0 | 0 | 0 |
| cluster delta area4:cluster delta vol4 | 0 | -0.04 | 0 | -0.04 | 0 | 0 | 0 |

Group LASSO models for clusters by applying a less conservative λ (lowest mean square error). β corresponds to the output coefficient. A value of 0 means that the predictors for the variable of interest were shrunk to 0. Sex and Age at baseline as covariates of no interest.

### Supplementary Figure 1

**a) PCA change in cortical thickness**

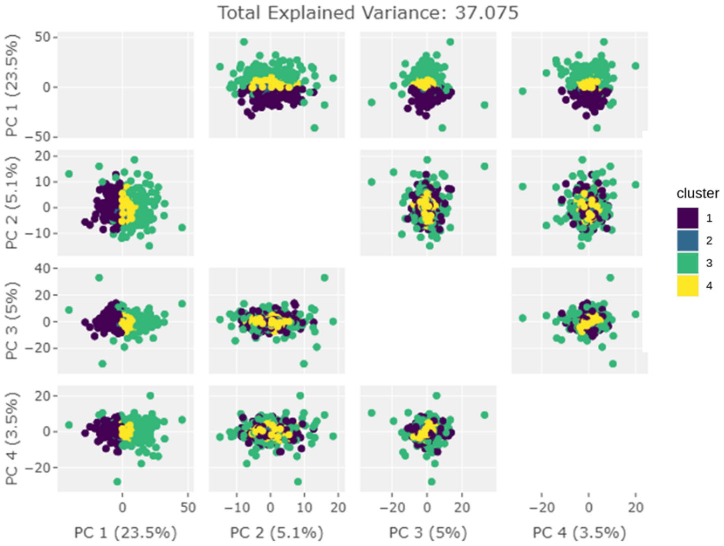


**b) PCA change in surface area**

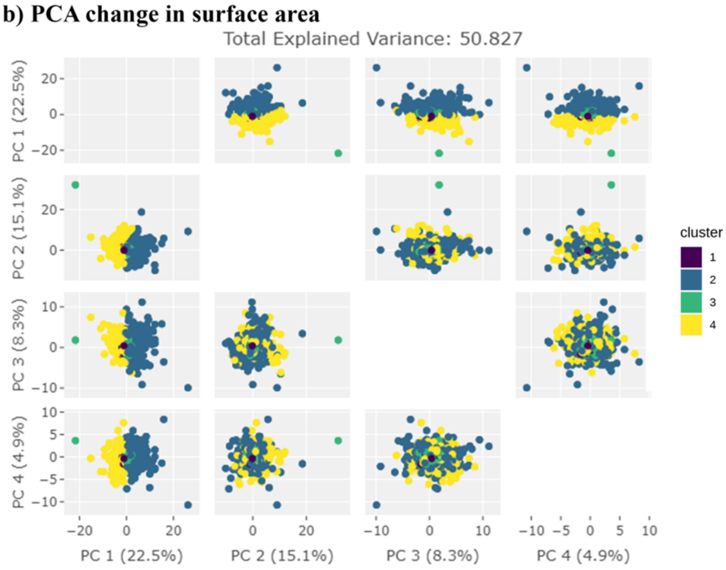


**c) PCA changes in subcortical volume**

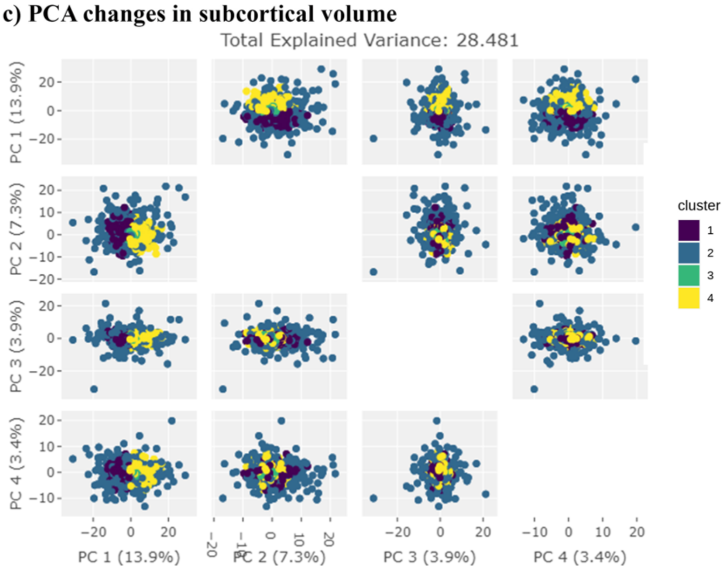

Principal component analysis onto clustering assignment solutions. A) PCA changes in cortical thickness, b) PCA changes in surface area, and c) PCA changes in subcortical volume. Each point represents an individual. The colour represents an individual's cluster assignment. For each modality, only the main four PCAs are shown. Note that different individuals organized into clusters can be observed only along the first PCA axis.
